# Genomic and Transcriptomic Signatures of SETD1A Disruption in Human Excitatory Neuron Development and Psychiatric Disease Risk

**DOI:** 10.1101/2025.03.26.645419

**Authors:** Zhixiong Sun, Huixiang Zhu, Xiaofu He, Bas Lendemeijer, Zanxu Wang, Jack Fan, Yan Sun, Zhiguo Zhang, Sander Markx, Steven A. Kushner, Bin Xu, Joseph A. Gogos

**Author notes:** Senior and corresponding authors (B.X), (J.A.G).

## Abstract

Genetic disruption of SETD1A markedly increases the risk for schizophrenia. To elucidate the underlying mechanisms, we generated isogenic organoid models of the developing human cerebral cortex harboring a *SETD1A* loss-of-function schizophrenia risk mutation. Employing chromatin profiling combined with RNA sequencing, we identified high-confidence SETD1A target genes, analyzed the impact of the mutation on SETD1A binding and transcriptional regulation and validated key findings with orthogonal approaches. Disruption of SETD1A function disturbs the finely tuned temporal gene expression in the excitatory neuron lineage, yielding an aberrant transcriptional program that compromises key regulatory and metabolic pathways essential for neurodevelopmental transitions. Although overall SETD1A binding remains unchanged in mutant neurons, we identified localized alterations in SETD1A binding that correlate with shifts in H3K4me3 levels and gene expression. These changes are enriched at enhancer regions, suggesting that enhancer-regulated genes are especially vulnerable to SETD1A reduction. Notably, target genes with enhancer-bound SETD1A are primarily linked to neuronal functions while those with promoter-bound SETD1A are enriched for basic cellular functions. By mapping the SETD1A binding landscape in excitatory neurons of the human fetal frontal cortex and integrating multimodal neuroimaging and genetic datasets, we demonstrate that the genomic context of SETD1A binding differentially correlates with macroscale brain organization and establish a link between SETD1A-bound enhancers, schizophrenia-associated brain alterations and genetic susceptibility. Our study advances our understanding of the role of SETD1A binding patterns in schizophrenia pathogenesis, offering insights that may guide future therapeutic strategies.

## Main

We have previously linked mutations in *SETD1A*, a histone methyltransferase, to risk for schizophrenia (SCZ), a disabling neuropsychiatric disorder defined by positive and negative symptoms as well as cognitive impairment^1,2^. This finding was confirmed in large-scale meta-analyses^3,4^ that established *SETD1A* as the gene currently with the most robust statistical support for association with SCZ risk. *SETD1A* is a widely expressed lysine methyltransferase which is best known as part of the Set/COMPASS complex that mediates mono-, di-, and tri-methylation of the lysine 4 on the histone H3 protein^5^. Several lines of evidence from human genetics and studies of postmortem human brain tissue suggest that histone H3 methylation is broadly relevant to the genetic risk and phenotypic variation of psychiatric and neurodevelopmental disorders^6–10^. In addition to SCZ, loss of function (LoF) and missense mutations in *SETD1A* have been found in individuals with disrupted speech development^11^ and early-onset epilepsy^12^ and described in the context of a neurodevelopmental syndrome whose core characteristics include global developmental delay (such as speech delay or motor delay) and/or intellectual disability, facial dysmorphisms, as well as behavior and psychiatric abnormalities^13^. Thus *SETD1A* mutations appear to increase neurodevelopmental vulnerability.

The study of *SETD1A* mutations in animal models has aided in our understanding of the affected cellular processes and signaling pathways in the adult brain that underlie the risk conferred by such mutations and provided important translational insights^14–16^. Nevertheless, if and how genetic disruption of *SETD1A* affects the unfolding of neurodevelopmental processes during human brain development remains largely unknown. Moreover, the genomic binding properties and targets of SETD1A in human forebrain neurons are largely unmapped. Brain organoids are amenable to prolonged culturing in a 3D-microenvironment that allows for an unforced generation of various cell types that mirror prenatal states of the human brain^17–19^ and provide a critical platform to dissect as well as integrate information, which would be otherwise difficult to assess *in vivo* within the developing human brain. To this end, here we introduce a *SETD1A* LoF risk mutation identified in patients with SCZ into a reference human induced pluripotent stem cell (hiPSC) line by CRISPR/Cas9 and generate dorsal forebrain organoids in order to determine the genomic binding features of SETD1A, identify SETD1A genomic targets and characterize the effect of *SETD1A* risk mutations on gene expression and neuronal development during early stages of forebrain development. Furthermore, by integrating chromatin profiling in the human fetal brain, transcriptomic analyses, and neuroimaging data, we link SETD1A-bound enhancers to schizophrenia-associated genetic and brain alterations, providing insights into disease mechanisms and potential therapeutic targets.

## Results

### Generation of hiPSC lines and forebrain organoids harboring a *SETD1A* LoF mutation

*SETD1A* LoF mutations confer a large increase in disease risk^1,3,4^, which provides an excellent start point for disease modeling. To this end, we adopted an isogenic experimental design and employed Clustered Regularly Interspaced Short Palindromic Repeats (CRISPR)/Cas9 genome engineering technology^20^ to introduce, in a well-characterized human control iPSC line (MH0159020)^21^, a *SETD1A* LoF mutation identified in patients with SCZ^1,2^ that creates a premature stop codon in exon 7 near the N-terminus of the protein [c.1272 delC, hereafter referred to as frameshift (FS)] (**Extended Data Fig.1a**). We obtained a sizable number of heterozygous mutants but no homozygous lines could be generated, suggesting that SETD1A plays an essential role in human iPSC survival, consistent with previous findings in mouse ESCs^22,23^. Two independent clones for wild type (WT) and FS genotypes were selected for further analysis. After confirming the successful introduction of the desired mutation (**Extended Data Fig. 1b**), we assessed its impact on SETD1A expression by immunoblotting and qRT-PCR, revealing that mutant iPSC lines exhibit approximately a 50% reduction in SETD1A mRNA and protein levels compared to WT (**Extended Data Fig. 1c**). Additionally, we verified that all iPSC lines retain their stemness through immunostaining for stem cell markers, maintain genomic stability via G-band karyotype analysis, and lack detectable off-target effects, as confirmed by Sanger sequencing of predicted off-target sites (**Extended Data Fig. 2**).

To investigate the effects exerted by the *SETD1A* mutation on human cortical development, we derived dorsal forebrain organoids from WT and mutant iPSC lines. *SETD1A* expression is low in iPSCs and greatly increases upon neuronal differentiation in organoids, consistent with its expression pattern in the developing human brain (**Fig. 1a**). Western blot assays confirmed a significant (p <0.05) reduction of SETD1A protein in 10-week-old mutant organoids, with no truncated SETD1A protein detected (**Fig. 1b**).

**Figure 1.**
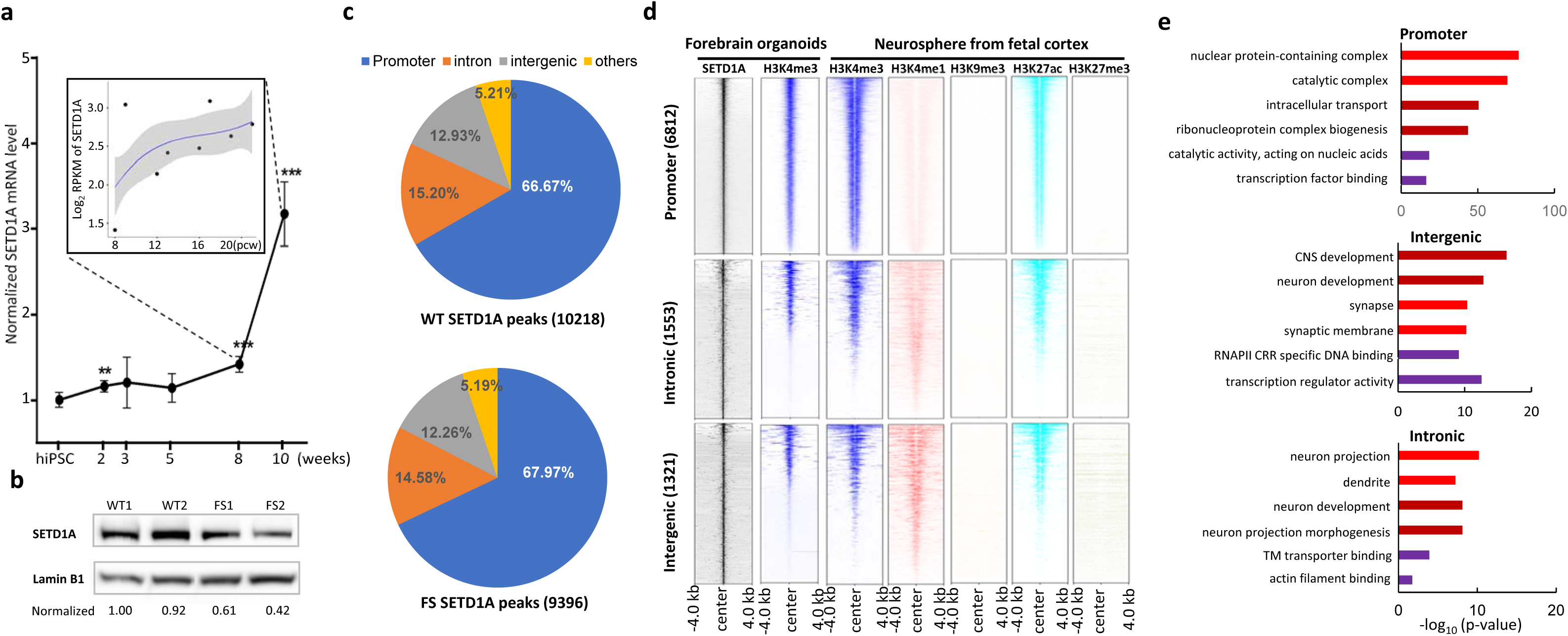
Genome-wide characterization of SETD1A binding sites in cortical EN derived from forebrain organoids. **a,** SETD1A expression profile during neuronal differentiation from hiPSCs to forebrain organoids. Image in box depicts the expression profile of SETD1A in human fetal dorsolateral prefrontal cortex. pcw: postconceptional weeks. RPKM: Reads Per Kilobase Million. **b,** Western blot analysis of SETD1A expression in WT and mutant forebrain organoids at DIV70, SETD1A protein levels were probed with mouse monoclonal antibody sc-515590 (Santa Cruz Biotechnology), quantified and normalized to a housekeeping gene encoding Lamin B1, and fold changes were shown as a relative value to WT1. Representative data of 2 independent experiments were shown. **c,** Genomic distribution of SETD1A peaks (n = 10218 in WT and n = 9396 in FS). **d,** Heatmap of histone marks (H3K4me3, H3K4me1, H3K9me3, H3K27ac and H3K27me3) centered on SETD1A peaks (n = 10218) across a ±4kb window. Histone modification profiles were from datasets of neurons sorted from forebrain organoids and extracted from ENCODE datasets from human fetal (17 weeks) cortex-derived neurospheres, and then compared to the SETD1A genomic occupancy map. SETD1A peaks were clustered in promoter (<1kb) (top, 6812 peaks), intronic (middle, 1553 peaks) and intergenic regions (bottom, 1321 peaks). **e,** GO analysis of genes with SETD1A binding sites in promoter, intronic and intergenic regions indicates that genes targeted by SETD1A at intronic and intergenic enhancer sites are more likely involved in neuronal-specific processes. CRR denotes cis-regulatory region.

### Genome-wide characterization of the SETD1A binding sites in cortical neurons of human forebrain organoids

To identify genomic binding sites of SETD1A in cortical neurons derived from organoids, we adopted the CUT&Tag approach, an ultra-sensitive ChIP-seq assay with high signal-to-noise ratio^24^, followed by deep sequencing (**Extended Data Fig.3a**) (see Methods). The specificity of the SETD1A antibody was confirmed by immunoprecipitation (IP) in HEK293T cells (**Extended Data Fig.3b**). Binding site identification was performed on sorted neurons from DIV70 organoids transduced with AAV1-hSYN-eGFP virus, which drives eGFP expression under the human SYNAPSIN 1 promoter exclusively labeling excitatory neurons (EN) while excluding progenitor neurons, as supported by human fetal brain data^25^ and further corroborated by the 2-month old cerebral organoid UMAP analyses^26^. The output sequence data were analyzed using the well-established ENCODE chip-seq pipeline2 for read alignment, quality control (QC, with a genome-wide Irreproducible Discovery Rate (IDR) < 0.05) and peak calling (MACS2, v.2.2.7.1). Experiments were conducted on 3 biological replicates from each individual clone, and 2 independent clones for each genotype. A merged peak list from three biological replicates for each clone was generated and a “consensus peak list” from the two independent clones of the same genotype was produced (see Methods). Employing an IgG-only negative control we filtered out 920 potential nonspecific pA-Tn5 enzyme insertion sites^27^ and the filtered “consensus peak lists” were used for downstream analysis. We detected 10218 high quality SETD1A peaks in WT and 9396 in mutant samples (**Fig. 3c**) (IDR <0.05 genome-wide, **Supplementary Table 1**).

To gain insights into the regulatory properties of the SETD1A binding sites, we first catalogued the genomic distribution of SETD1A peaks according to their positions in cis-regulatory elements. In WT neurons, we found that the majority of SETD1A binding sites (66.67%, 6812) are located in promoters [within 1kb of the transcription start sites (TSS)], followed by introns (15.20%, 1553) and intergenic regions (12.93%, 1321). A minority of binding sites (5.21%, 532) are located in other regions such as exons, 3’ UTR, 5’ UTR and transcription termination sites (TTS) (**Fig. 3c**). Similarly, in mutant neurons, 67.97% (6386) of SETD1A binding sites are located in promoters, followed by introns (14.58%, 1370) and intergenic regions (12.26%, 1152). Comparison of the genomic distribution pattern of SETD1A peaks in WT and mutant samples indicates that the *SETD1A* haploinsufficiency does not lead to a global redistribution of its cis-elements occupancy (**Fig. 3c**).

We examined the genomic distribution of H3K4me3 peaks using CUT&Tag and identified 14496 peaks in WT neurons and 14687 peaks in mutant neurons (**Extended Data Fig. 3c**, IDR <0.05 genome-wide, **Supplementary Table 1**). Consistent with H3K4me3 being a well-established promoter mark, these peaks are predominantly located in promoter regions. Furthermore, the peaks exhibited high enrichment in SETD1A-bound promoter regions, in line with SETD1A’s known H3K4me3 methyltransferase activity (**Fig. 3d)**. Comparison of the global H3K4me3 binding profiles in WT and mutant samples did not reveal any widespread redistribution of binding sites (**Extended Data Fig. 3c**).

To evaluate the fidelity of the identified SETD1A peaks in organoid derived cortical neurons, we extracted publicly available histone modification profiles (including H3K4me1, H3K4me3, H3K9me3, H3K27ac and H3K27me3) from ENCODE datasets produced from human embryonic (17 week) cortex-derived neurospheres (**Supplementary Table 2**) and conducted a detailed comparison of our SETD1A genomic occupancy map against the ENCODE catalogue. SETD1A catalyzes the tri-methylation of H3K4 in promoter regions and mono-methylation of H3K4 in enhancers^28^. Consistently, we observed a higher than expected by chance overlap between SETD1A binding sites and histone markers for promoters (H3K4me3, and H3K27ac) and enhancers (H3K4me1, and H3K27ac) (p = 1×10^-5^, Z = 140.0 for H3K4m1; p = 1×10^-5^, Z = 436.2 for H3K4m3; p = 1×10^-5^, Z = 332.7 for H3K27ac by RegioneR). By contrast, SETD1A binding sites exhibited either a lower overlap with or depletion in other histone markers (p = 1×10^-5^, Z = - 7.2 for H3K9m3 and p = 1×10^-5^, Z = 4.3 for H3K27m3) from the ENCODE repository (**Extended Data Fig. 3d**). We obtained similar results using additional ENCODE histone marker datasets from human embryonic (17 week) brain and human H9 embryonic stem cell-derived neurons (**Extended Data Fig. 4a**). To control against biases arising from our classification system, we also classified SETD1A binding sites according to a recently published machine learning-based annotation of cis-regulatory regions to promoter-like elements (PLS), proximal enhancer-like elements (pELS) and distal enhancer like elements (dELS)^29^. We found that 6523 (63.84%) of SETD1A peaks are located at promoter-like elements (PLS) overlapping with H3K4me3 and H3K27ac peaks while 1548 (15.15%) of SETD1A peaks are located at proximal enhancer-like elements (pELS), and 1725 (16.88%) of SETD1A peaks are located at distal enhancer-like elements (dELS) overlapping with H3K4me1 and H3K27ac peaks (**Extended Data Fig. 4b**).

Given that the histone code can play a role in higher-order folding of chromatin^30–32^, we also explored whether there is an enrichment of higher-order chromatin contacts among SETD1A binding sites. To this end, we analyzed HiC-seq data from the cortical and subcortical plate of human fetal cerebral cortex^33^ (**Supplementary Table 2**) using a permutation test (N=50,000 permutations implemented in the RegioneR program) and found a highly significant overlap between SETD1A binding sites and chromatin contacts (*P* = 2×10^-5^, Z=10.6) (**Extended Data Fig.4c**). This finding indicates that SETD1A may modulate 3D chromatin architecture and regulate gene expression at both local and distal cis-regulatory elements.

Finally, *de novo* motif analysis using hypergeometric optimization of motif enrichment (HOMER) at SETD1A peaks, to identify auxiliary transcription factor (TF) recruiting SETD1A to promoters and enhancers, revealed 18 de novo motifs as significantly enriched within the SETD1A binding sites (FDR p < 10^-4^, **Extended Data Fig.4d, Supplementary Table 3**) and collectively covering 92% (9409 of 10218) of the total SETD1A-bound regions. The interactions between SETD1A and these auxiliary TFs likely mediate the effects of SETD1A on cell type and developmental stage specific gene expression.

### Functional enrichment among SETD1A genomic targets

We identified a total of 8187 target genes encompassing the SETD1A peaks detected in WT neurons. We conducted Gene Ontology (GO) analysis to dissect potential unique functions served by SETD1A binding at distinct cis-regulatory regions. Target genes with promoter SETD1A binding sites (n=5981) show the highest enrichment scores for terms related to “nuclear protein-containing complex”, “intracellular transport” and “transcription factor binding”. By contrast, target genes with the intronic binding sites (n = 1349) are strongly associated with terms related to “neuron development”, “neuron projection”, while genes with intergenic binding sites (n=1679) are enriched for terms related to “synaptic membrane” (**Fig. 1e**). Thus, genes targeted by SETD1A at the intronic and intergenic enhancer sites are more likely involved in key neuronal-specific process. These findings are robust across annotation methods, as machine learning-based classification of SETD1A binding sites similarly showed that genes overlapping PLS are enriched for core transcriptional functions, while those overlapping pELS and dELS are primarily linked to neuronal processes (**Supplementary Table 2**).

Further refined functional annotation *via* mapping of intronic and intergenic SETD1A-bound target genes associated with the “neuron development” GO term (**Supplementary Table 3**) to the Search Tool for Retrieval of Interacting Genes (STRING) revealed a large number of genes connected by a PPI network highly enriched for biological processes related to “Generation of neurons” and “Neuron projection”) (**Extended Data Fig. 5a**). Supporting the enrichment of neuronal functions in SETD1A-bound intronic and intergenic regions, functional annotation revealed that target genes with an RFX3 motif in these regions are significantly enriched for “neuron development” terms. In contrast, genes with an RFX3 motif in SETD1A-bound promoters are enriched for “cilium organization,” a well-established role of RFX3^34^, underscoring the distinct functions of enhancer versus promoter binding sites (**Extended Data Fig. 5b).**

We found that 2190 (26.8%) of the target genes carry two or more SETD1A binding sites (**Supplementary Table 3**). As expected, there is a positive correlation between the number of SETD1A binding sites and the length of the associated target genes, with genes harboring two or more binding sites being significantly longer than target genes with one binding site (**Extended Data Fig. 5c-d**) and displaying a higher abundance of SETD1A-bound enhancer elements (**Extended Data Fig. 5e**).

### Transcriptional dysregulation induced by the SETD1A LoF mutation in cortical neurons

To identify transcriptomic changes resulting from the SETD1A LoF mutation, we conducted bulk RNA-seq analysis and parallel genomic assessment of neurons derived from a different batch of WT and FS mutant organoids. There was a very good agreement between batches in the identity, genomic distribution and regulatory properties of SETD1A binding sites (**Extended Data Fig. 6**, **Supplementary Table 4**). PCA analysis of RNA-seq data shows that WT and mutant groups are transcriptomically distinct (**Extended Data Fig. 7a**). Bulk RNA-seq deconvolution, leveraging RNA-seq data from cells immunopurified from human brain tissues^35^, revealed that 99.89%–100% of cells in both WT and mutant samples were neurons, confirming the homogeneity of the cell populations (**Supplementary Table 5**). Notably, despite the reduction of SETD1A protein in 10 weeks old mutant organoids (**Fig. 1b**), RNA-seq analysis indicated the presence of mutant transcripts and confirmatory qRT-PCR did not reveal a significant decrease in SETD1A transcript levels suggesting that, unlike in iPSCs, mutant SETD1A transcripts are not efficiently targeted for degradation in developing neurons (**Extended Data Fig. 7b**).

We detected 806 differentially expressed genes (DEGs) in mutant neurons (DESeq2, padj <5%) with 532 (66.0%) up-regulated and 274 (34.0%) down-regulated (**Fig. 2a, Supplementary Table 5**). DEGs are enriched in GO terms related to “synaptic membrane”, “glutamatergic synapse” and “neuron projection” (**Fig. 2b**). We identified 421 DEGs (52.23%) that carry one or more SETD1A binding sites in their regulatory elements, likely representing primary rather than secondary transcriptional alterations. Functional annotation of these SETD1A target DEGs further revealed enrichment in GO terms relevant to “neuron development”, “neuron projection”, “synaptic membrane” as well as “voltage gated potassium channel activity” (**Fig. 2c** and **Extended Data Fig. 7c**). Transcriptional dysregulation of genes involved in neuronal projections and synaptic function predicts alterations in neurite outgrowth, synaptic transmission and plasticity, consistent with findings in human iPSC-derived neurons carrying heterozygous disruptions of SETD1A^36–38^. Downregulation of genes associated with “voltage-gated potassium channel activity”, crucial for action potential (AP) dynamics, predict changes in the intrinsic excitability of mutant neurons. We conducted whole-cell patch-clamp recordings on neurons differentiated from purified populations of NPCs and found that mutant neurons exhibited a lower AP threshold (p < 0.001), a greater AP amplitude (p < 0.001) and a higher maximum firing rate (p < 0.001). Additionally, the AP half-width was significantly narrower (p < 0.001), reflecting faster repolarization kinetics. These findings were consistent across two independent WT and mutant lines (**Fig. 2d** and **Extended Data Fig. 7d**). Changes in resting membrane potential, input resistance, and capacitance were less pronounced (**Extended Data Fig. 7e**), suggesting that the primary alterations in excitability are linked to active membrane properties.

**Figure 2.**
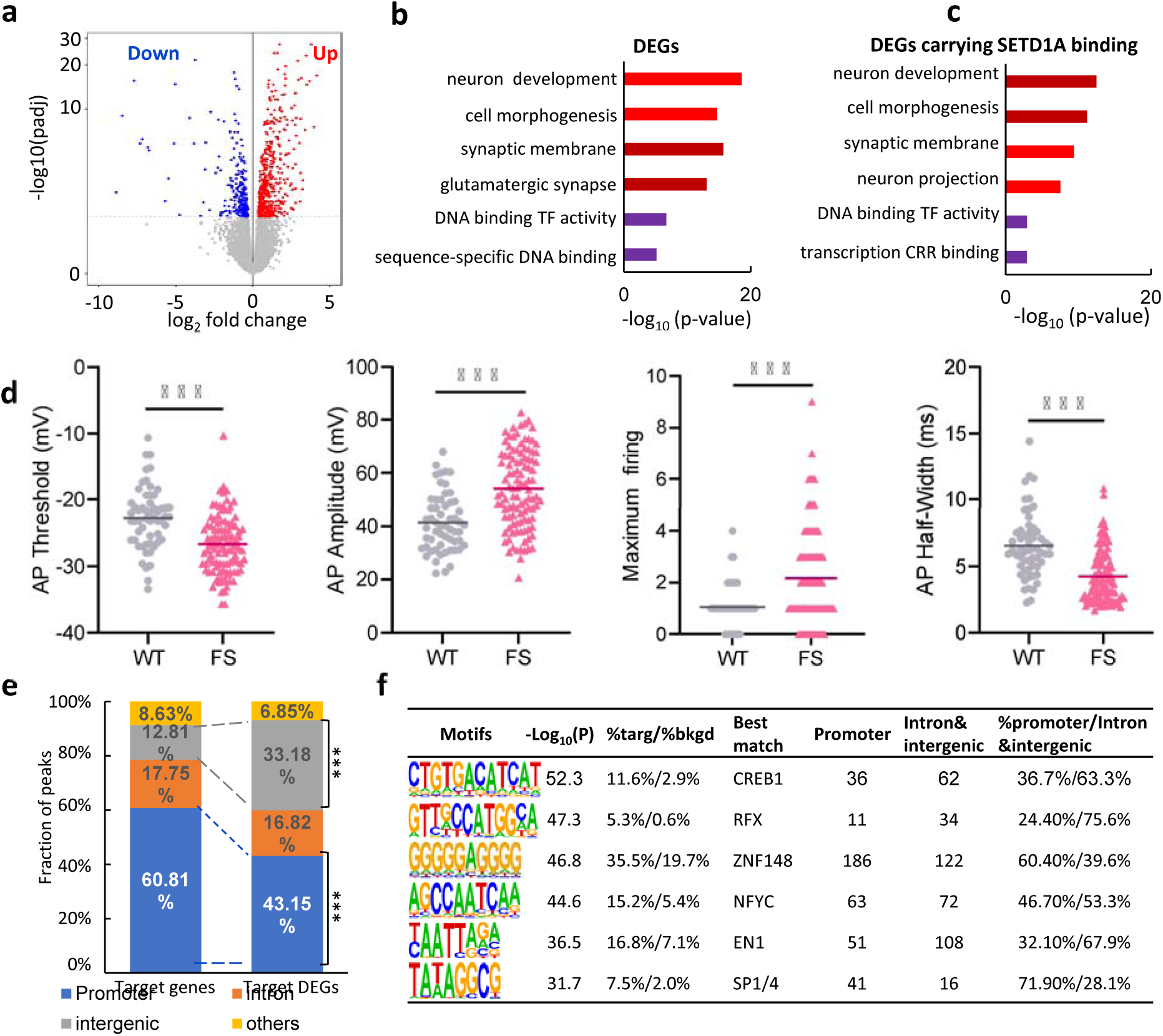
Transcriptional alterations induced by the *SETD1A* LoF mutation in developing human cortical neurons. **a**, Volcano plot of RNA-seq result showing DEGs in sorted neurons from DIV70 mutant organoids compared to WT organoids. **b-c,** GO enrichment analysis of DEGs (**b**) and SETD1A target DEGs (**c**). **d,** Active electrophysiological properties in WT and mutant neurons. Quantification of action potential (AP) threshold, AP amplitude, Maximum firing and AP half width were performed, n>50 neurons were examined. **e,** Genomic distribution of SETD1A peaks overlapping with DEGs. Permutation analysis indicates that compared to SETD1A binding sites as a whole, SETD1A peaks overlapping with DEGs are more frequent in intronic and intergenic regions, ***p < 0.001. **f,** De novo TF motif analysis of SETD1A binding sites residing within DEGs.

Notably, among the 674 SETD1A peaks encompassed within target DEGs, 291 (43.18%) are located in promoters, 114 (16.91%) in introns, 223 (33.09%) in intergenic regions and 46 (6.82%) in other regions. Thus, compared to SETD1A binding sites as a whole, SETD1A peaks overlapping with target DEGs are more frequent in intronic and intergenic regions (50% versus 28.13%), and less frequent in promoter regions (43.15% versus 60.81%). This shift in the genomic distribution is statistically significant (**Fig. 2e**, empirical *P* < 1×10^-3^, **Extended Data Fig. 7f**) and is observed among target DEGs independently of the number of their SETD1A binding sites (**Extended Data Fig. 7g**). Collectively, these data suggest that genes with SETD1A bound enhancers may be more susceptible to the effects of *SETD1A* haploinsufficiency. Consistently, genes with multiple SETD1A binding sites and a higher abundance of SETD1A-bound enhancer elements (**Extended Data Fig. 5c**) are significantly enriched among target DEGs as compared to SETD1A targets as a whole (**Extended Data Fig. 7h**).

*De novo* motif analysis of SETD1A binding sites residing within DEGs identified 6 significantly enriched *de novo* motifs. This list includes an RFX consensus motif (**Fig. 2f, Supplementary Table 5**), one of the top-ranked motifs enriched within the entire set of SETD1A binding sites. Approximately 70% of the SETD1A peaks carrying an RFX motif are located in intronic and intergenic regions, an ∼1.5-fold enrichment over the SETD1A binding sites as a whole (44.61%) (**Extended Data Fig. 7i**). SETD1A target DEGs carrying an RFX motif are enriched for GO terms related to neuronal development and include genes such as CAMK2, PICK1, GAP43 and NTRK2. These observations support to the notion that RFX family members play an important role in mediating the effects of SETD1A on gene expression during early human development of cortical neurons. Interestingly, this list also includes a consensus motif for SP4^39^, a top-ranked SCZ risk gene supported by both exome sequencing^4^ and GWAS^40^ data. SETD1A target DEGs carrying an SP4 motif include genes such as FOXG1, NTRK2, SLIT2 and PICK1 and are also enriched for GO terms related to neuronal development, indicating a potential interaction of SETD1A and SP4 in regulating gene transcription in developing human cortical neurons.

### Differential SETD1A binding sites in mutant neurons are enriched in intronic and intergenic regions

Reduction of SETD1A levels in mutant neurons does not lead to global redistribution of SETD1A binding but may affect local binding occupancy. To ensure the reliable identification of differential binding sites (DBSs) between WT and mutant neurons, we incorporated *E. coli* tracer DNA as a spike-in for normalization and performed differential SETD1A binding analysis using Linnorm^41^. We identified 351 SETD1A-bound peaks displaying differential binding strength (**Fig. 3a**, **Supplementary Table 6**, Linnorm with FDR adjusted p value = 0.1, see methods). 158 DBSs display diminished SETD1A occupancy and 193 DBSs display enhanced SETD1A occupancy in mutant samples. The latter finding is unexpected and may be related to mutation-associated perturbations in auxiliary TF factor expression or local chromatin topology that alter the accessibility of regulatory regions in *SETD1A*-deficient cells.

**Figure 3.**
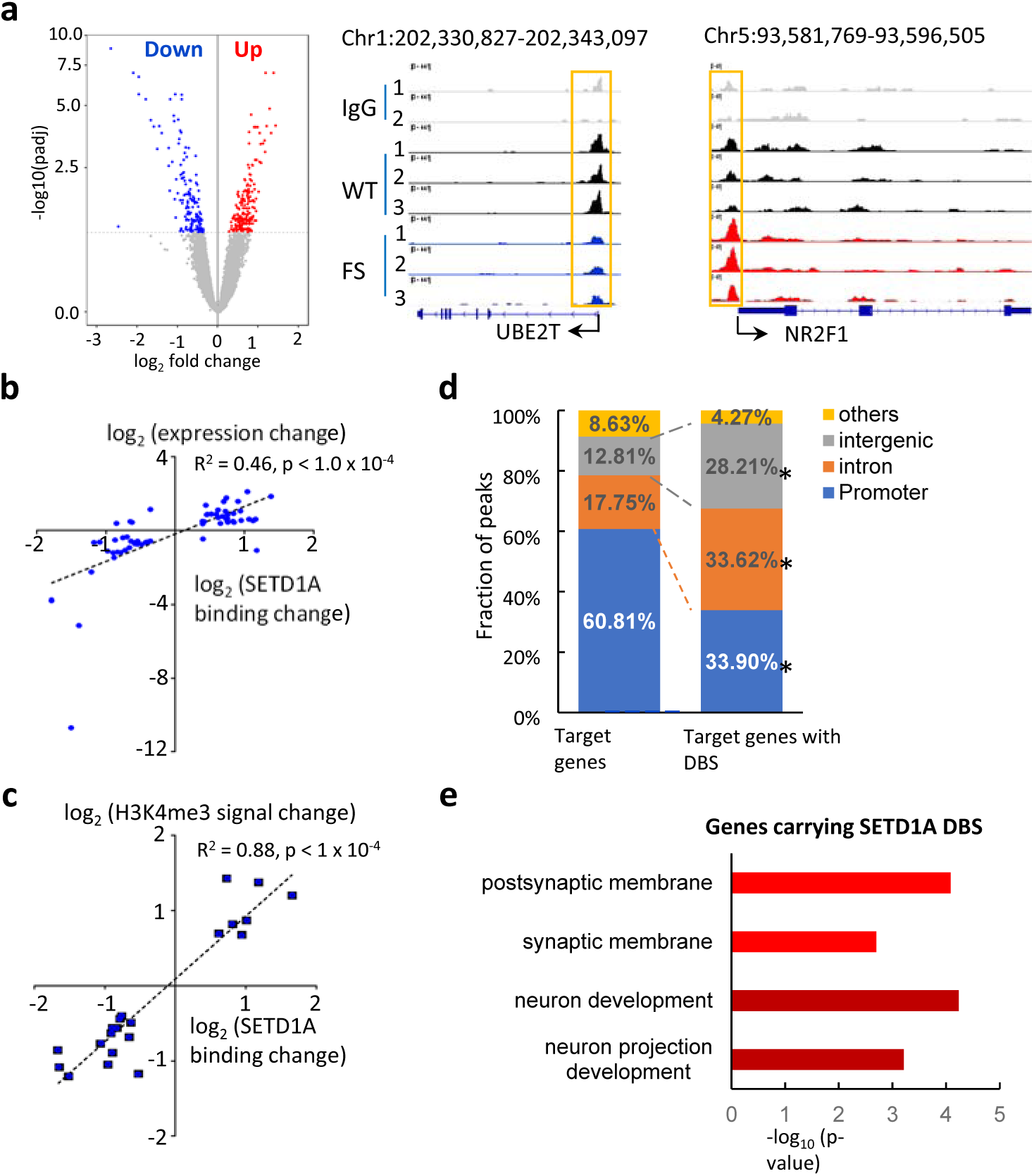
Characterization of Differential SETD1A Binding Sites in mutant cortical neurons. **a**, Volcano plot of differential SETD1A binding analysis result showing DBS in sorted neurons from DIV70 mutant organoids compared to WT organoids (left). Representative DBS loci are also shown (middle, right). Bigwig files are displayed in Integrated Genomics Viewer and green boxes highlight the actual DBS loci. Numbers 1-3 denote the three biological replicates of the CUT&Tag analysis. **b,** Correlation analysis between SETD1A DBS and DEGs found a positive correlation between the direction of DEG expression and SETD1A occupancy changes in associated DBS; x axis depicts log2 binding occupancy fold change, y axis depicts log2 gene expression fold change. **c,** Correlation analysis between H3K4me3 and SETD1A DBS found a positive correlation between the direction of H3K4me3 signal and SETD1A occupancy changes; x axis depicts log2 SETD1A binding occupancy fold change, y axis depicts log2 H3K4me3 signal fold change. **d,** Genomic distribution of DBS. Permutation analysis indicates that compared to SETD1A binding sites as a whole, DBSs show a higher combined localization in intronic and intergenic as compared to promoter regions. *p < 0.05, unpaired Student’s t test. **e,** GO term enrichment analysis of genes carrying SETD1A DBS.

To elucidate the functional implications of this unexpected finding, we conducted two follow-up analyses and discovered compelling evidence of its impact on gene regulation. First, we identified 61 target DEGs in sorted neurons that overlap with DBS (**Supplementary Table 6**) and found a significant positive correlation between the direction of gene expression and SETD1A occupancy changes in associated DBS (R^2^ = 0.46, *p* < 1.0 x 10^-4^) (**Fig. 3b**) providing a strong functional link between the observed enhanced SETD1A binding and SETD1A target gene upregulation in the mutant neurons. Second, we examined whether local alterations in SETD1A occupancy were accompanied by corresponding changes in H3K4me3 levels. Differential binding analysis between WT and mutant samples identified 97 sites with differential H3K4me3 signal, indicating that the SETD1A mutation does not lead to widespread H3K4me3 redistribution. Among these, 21 sites (16 in promoters and 5 in enhancers) overlapped with SETD1A DBS, and at these overlapping regions, changes in H3K4me3 signal were strongly positively correlated with SETD1A occupancy (R² = 0.88, p < 1 × 10⁻⁴, **Fig. 3c**), with increased SETD1A occupancy corresponding to elevated H3K4me3 signal and vice versa. Among the 21 overlapping sites, 11 (52.4%) were linked to DEGs, all showing a consistent positive correlation between SETD1A occupancy, H3K4me3 level, and target gene expression changes(**Supplementary Table 6, Extended Data Fig. 8a**, R² = 0.56, p = 0.0082), suggesting that, at specific sites, SETD1A mutation alters transcription by modulating H3K4me3 levels.

Compared to SETD1A binding sites as a whole, DBSs show a higher combined localization in intronic (33.62%) and intergenic (28.21%) regions as compared to promoters (33.90%) (**Fig. 3d**). Because SETD1A peaks in intronic and intergenic regions are significantly shorter than those in promoters, we examined the distribution of DBSs across SETD1A peaks of varying lengths and found that shorter peaks, which may be more susceptible to displaying differences in our assay, did not disproportionately contribute to DBS frequency. By contrast, we observed a modest increase of DBS in SETD1A peaks with longer length, thus ruling out the possibility that DBS enrichment in enhancer regions arises from a bias associated with shorter peak lengths (**Extended Data Fig. 8b**). The shift in the genomic distribution of DBSs is statistically significant (**Extended Data Fig. 8c-e**, empirical *P* < 1×10^-3^ for promoter, intronic and intergenic regions, respectively) and indicates that SETD1A binding within intronic and intergenic enhancers is particularly vulnerable to the effects of SETD1A mutations relatively to promoter binding. In line with the notion that, for a subset of SETD1A binding sites, SETD1A mutation alters transcription *via* H3K4me3 modulation, we observed a similar enrichment in enhancer regions of sites with differential H3K4me3 signal (**Extended Data Fig. 8f**). Functional annotation found that target genes carrying SETD1A DBS sites display enrichment in “neuron development” and “synaptic membrane” (**Fig. 3e, Supplementary Table 6**).

### SETD1A mutation alters early developmental trajectories of Ens

Our analysis of SETD1A target genes and of the transcriptomic changes induced by a *SETD1A* LoF mutation identified robust molecular signatures indicative of curtailed development and differentiation. To examine whether these molecular alterations are reflected in cell population alterations and examine *in situ* genotypic differences in transcriptomic and regulatory profiles during neurogenesis and neuronal differentiation, we used single cell RNA sequencing (scRNA-seq) to profile cell type composition and molecular signatures of organoids derived from mutant and WT iPSC lines at DIV70, when neurogenesis is ongoing. A high viability (>90%) single cell suspension from 3 organoids per line was obtained for scRNA-seq. Our quality control metrics indicated that scRNA-seq data were uniform across the samples, with low counts of mitochondrial and ribosomal genes detected and comparable count metrics (**Extended Data Fig. 9a-c**). A total of 16,541 single cells from DIV70 organoids were analyzed following quality control and filtering. To annotate the cells in our scRNA-seq dataset, we adapted published datasets that classify the signature transcriptomic profiles of the cell-types within the human fetal forebrain at various stages of development^25^. We found that majority of cell types shown in the developing human brain are represented in our dataset (**Fig. 4a and Supplementary Table 7**).

**Figure 4.**
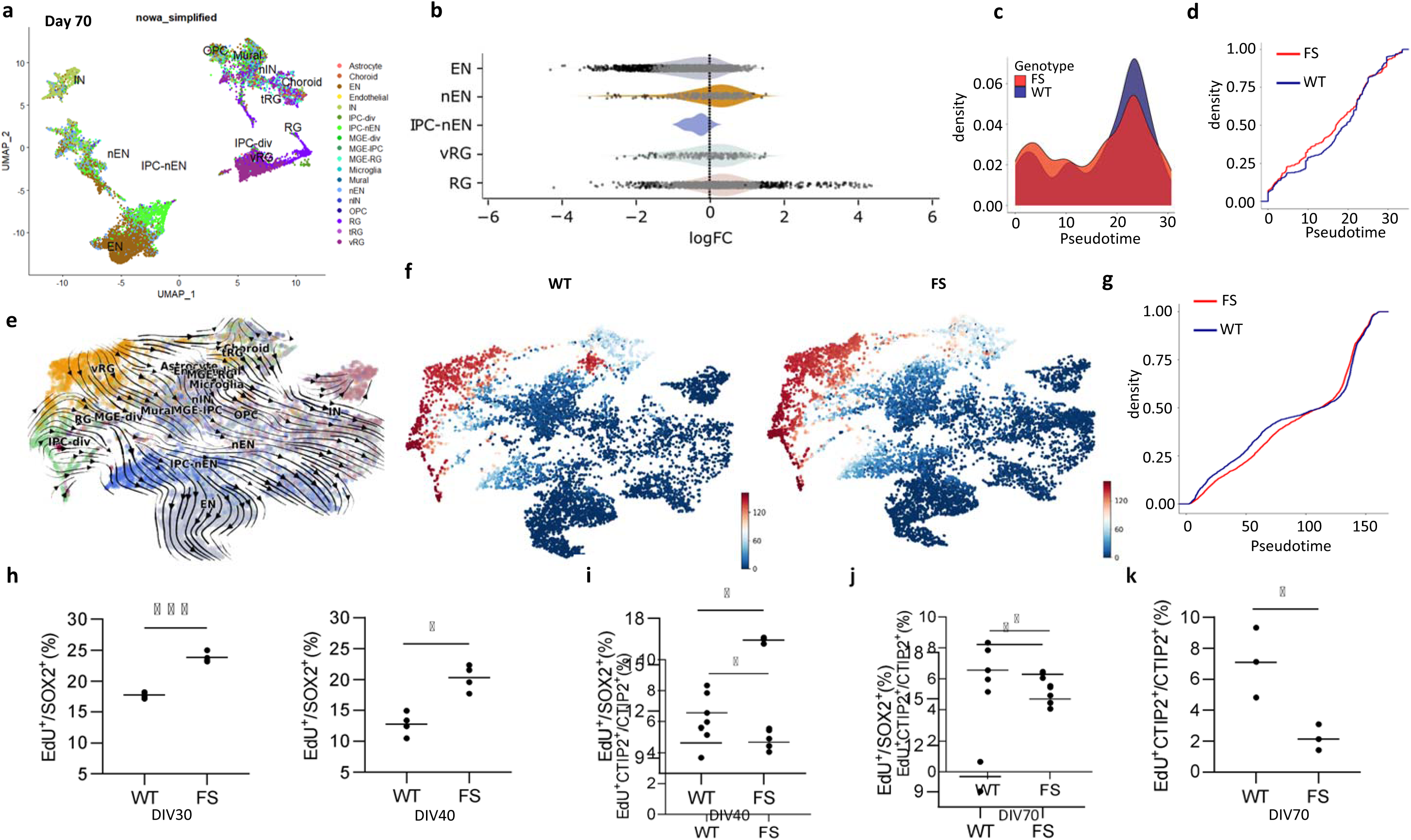
*SETD1A* mutation leads to asynchronous developmental trajectories in DIV70 forebrain organoids. **a**, Annotation of cell types in the dataset with reference to human fetal brain^36^. Most of cell types in the developing human fetal cortex are identified in DIV70 organoids. UMAP color coded to highlight distribution of annotated cell types in DIV70 organoids. **b**, Beeswarm plot depicting log fold change (Mutant/WT) in neighborhoods containing cells from the same cell type at DIV70. Mutant organoids have higher fraction of neural progenitor cells (RG) and lower fraction of differentiated ENs compared to WT organoids (spatialFDR <0.1). **c,** Pseudotime trajectory of EN lineage calculated with Monocle3. Ridge plot depicting distribution of cells in mutant and WT EN lineage at DIV70. Mutant cell distribution is shifted towards early phase of the trajectory represented by progenitor cells while WT cell distribution is shifted towards the terminal phase of the trajectory represented by more mature ENs. **d,** K-S test indicating a significant difference between the two distributions (P = 1×10^-5^). ECDF: Empirical Cumulative Distribution Function. **e,** UMAP plots of the RNA velocity embedding streams of mutant and WT organoid scRNA-seq datasets at DIV70 from neural progenitors (RG) to terminal neurons. **f,** UMAP plots colored by absorption probabilities in WT (left) and mutant (right) DIV70 EN lineage. Progenitors in WT organoids show high absorption probabilities while progenitors in mutant organoids show low absorption probabilities. **g,** K-S test indicating a significant longer mean time of absorption in mutant organoids (P = 9×10^-6^). **h,** Quantitative flow cytometry analysis revealing that mutant organoids exhibit a significantly higher proliferation rate (EdU⁺/SOX2⁺ cells) than WT organoids at DIV30 (left) and DIV40 (right). **i,** Quantitative flow cytometry analysis of an additional independent pair of organoids revealed that mutant organoids exhibit a significantly higher proliferation rate (EdU⁺/SOX2⁺ cells) than WT organoids at DIV40. **j,** Quantitative flow cytometry analysis revealing a significantly lower neurogenesis rate (EdU⁺CTIP2⁺/CTIP2⁺) in mutant organoids at DIV70 compared to WT organoids. **k,** Quantitative flow cytometry analysis of an additional independent pair of organoids revealed a lower neurogenesis rate (EdU⁺CTIP2⁺/CTIP2⁺) in mutant organoids at DIV70 compared to WT organoids. *** p < 0.001, ** p < 0.01, * p < 0.05, unpaired Student’s t test.

Quantification of the various cell types by Milo^42^ revealed a significant lower proportion of the major differentiated EN and higher proportion of neural progenitors (RG, vRG, and IPC-nEN) in mutant organoids (FDR < 0.1, **Fig. 4b**). We determined if cell composition changes result from aberrant neural lineage developmental trajectories using two independent approaches. First, we used Monocle3 to calculate the distribution of EN lineage cells along the pseudotime trajectory, as defined by transcriptomic changes, and map the developmental status of all cells along the pseudotime (**Fig. 4c and Extended Data Fig. 9d**). Comparison between mutant and WT organoids, revealed an increased distribution of mutant cells in the early phase of the pseudotime trajectory (progenitor states) and a decreased distribution towards the terminal phase of the trajectory (mature neuron states) (p < 0.0005, two-sided K-S test; **Fig. 4d**) supporting a delay in progenitor cell transition in mutant organoids. To corroborate with a complementary approach, we conducted single cell velocity analysis on the scRNA-seq dataset to calculate RNA velocity, a measure of the directed dynamics of expressed genes that leverages mRNA splicing kinetics using scVelo (**Fig. 4e**). We then used RNA velocity information and the CellRank pipeline to compute absorption probability, which estimates how fast each cell reaches the EN state, the endpoint of the lineage (**Fig. 4f**). A K-S test on the distribution of the absorption probability scores by genotype indicated that progenitor cells in mutant organoids take a longer time to reach the terminal state (P< 9×10^-6^, two-sided K-S test; **Fig. 4g**).

To experimentally validate the observations from our scRNA-seq analysis, we used fluorescent activated cell sorting (FACS) of cell suspensions obtained from mutant and WT organoids generated from two independent pairs of WT and mutant iPSC lines. Since the proportion of cycling NPCs is a small fraction of total cells in DIV70, we conducted cell cycle analysis at DIV30-40 organoids when there is a preponderance of actively cycling cells. We pulsed organoids with 5-ethynyl-2′-deoxyuridine (EdU) for 4 h to pulse-label proliferating cells in S phase, co-immunostained with SOX2 to label NPCs and quantitatively assessed proliferation with flow cytometry (**Extended Data Fig. 9e**). In agreement with the scRNA-seq analysis, experiments using independent WT and mutant iPSC lines demonstrated a significant increase in proportion of EdU^+^ cells among all SOX2^+^ in mutant organoids, indicating a significant expansion of proliferating progenitor cells (**Fig. 4h, i**). We also used FACS-based analysis to experimentally validate the reduced number of neurons in the scRNA-seq dataset. We analyzed sorted newborn neurons marked by EdU^+^CTIP2^+^ at DIV70 and confirmed a significant reduction in their proportion among all CTIP2^+^ labeled neurons, a finding observed across independent WT and mutant iPSC lines (**Fig. 4j, k**). Taken together, our findings suggest that the observed genotypic differences in transcriptomic and regulatory profiles have important consequences for the development of mutant NPCs.

### Gene expression alterations underlying aberrant EN early developmental trajectories

To determine the molecular mechanisms underlying the aberrant unfolding of neurodevelopment, we first used MAST software to search for DEGs in organoid neural progenitors and ENs (padj <5%, **Supplementary Table 7**). Analysis of all SETD1A target DEGs in ENs (**Supplementary Table 7)** found, in agreement with our bulk RNA-seq results in sorted neurons, a consistent positive correlation between the direction of gene expression and SETD1A/H3K4me3 signal changes in associated DBS (R^2^ ranging from 0.47 to 0.72, **Extended Data Fig. 10**).

To identify DEGs critical for differentiation, we used CellRank^43^ to identify 1624 putative lineage driver genes (genes whose expression are systematically higher or lower in cells biased towards a particular fate within the WT EN lineage, q value < 0.05) and found that 112 of them overlap with DEGs in ENs (**Supplementary Table 7**). Among these lineage driver genes affected by the SETD1A LoF mutation there are 83 SETD1A target genes (**Fig. 5a**). The pattern of expression of these target genes is affected by SETD1A disruption with many of them displaying marked disparities between WT and mutant organoids in their dynamic temporal expression profile (**Extended Data Fig. 11a**). Comparison to our bulk RNA-seq results identified 20 driver genes (13 targets and 7 non-targets) that display consistent changes in both scRNA-seq and bulk RNA-seq datasets (in terms of both p value and fold-change direction, **Supplementary Table 7)** as high confidence representative examples of individual dysregulated driver genes (**Fig. 5b** and **Extended Data Fig. 11b**). The majority of these genes (16/20) are upregulated in ENs, primarily due to failure in downregulating their expression during transition to more mature cellular states. Notably, among the 13 SETD1A target driver genes, 6 genes (NR2F1, NR2F2, ZFHX3, THSD7A, UBE2T and VIM, **Fig. 5b**) contain a DBS, a marked enrichment over the SETD1A target gene set as a whole (46.2% versus 2.5%). Five out of these 6 target genes are upregulated and 4 of them display enhanced SETD1A binding in mutant neurons (**Supplementary Table 7)**. Among dysregulated SETD1A target driver genes, notable examples include (i) TFs such as NR2F1 and NR2F2, which regulate neural progenitor fate^44^ and enhance neurogenesis when knocked down, showing upregulation in mutant organoids only at intermediate or late developmental stages but not in early progenitors. (ii) Metabolic enzymes, such as LDHA, whose downregulation marks the transition from aerobic glycolysis to neuronal oxidative phosphorylation as neurons develop from neural progenitor cells^45^, which remains elevated in mutant organoids at later stages. (iii) Membrane proteins such as NRP2, a member of the neuropilin family involved in axon guidance and elongation^46^ and THSD7A, a membrane-associated N-glycoprotein with pleiotropic effects implicated to bipolar disorder (BPD) genetic risk^47^, which are downregulated and upregulated, respectively, in mutant organoids at late time points. Among non-target dysregulated driver genes, notable examples include: (i) RBFOX1, a splicing regulator critical for neurodevelopment^48^ which is markedly upregulated at late developmental stages. (ii) PCDH10, a δ2-protocadherin family member involved in cell adhesion, which shows increased expression at intermediate and late stages. and (iii) ACAT2, which encodes cytosolic acetoacetyl-CoA thiolase, an enzyme involved in lipid metabolism, and it markedly downregulated at late stages (**Fig. 5b**). These findings suggest that disrupted SETD1A binding in mutant NPCs perturbs the temporal regulation of key genes, impacting both developmental and metabolic processes.

**Figure 5.**
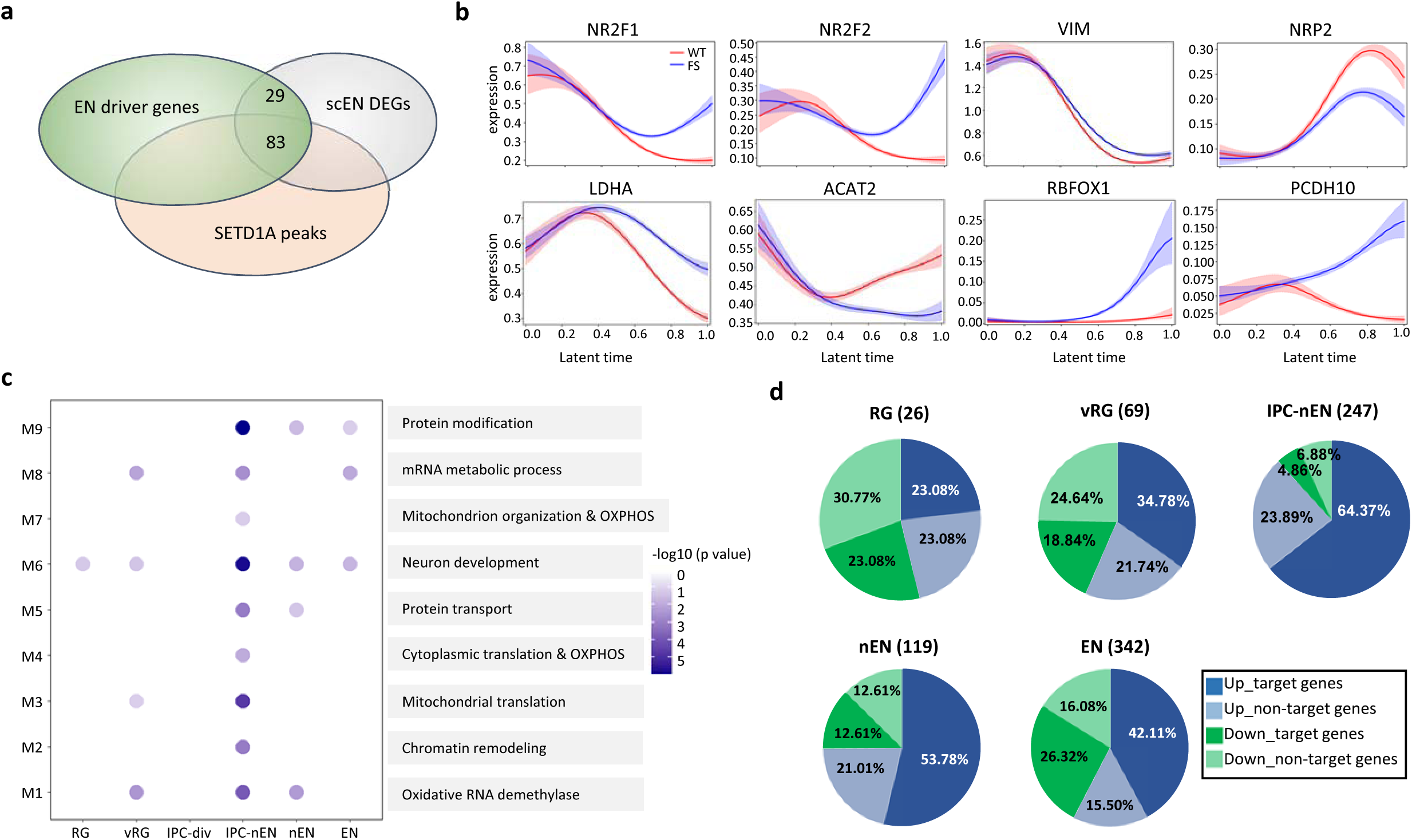
Aberrant gene expression underlying altered EN developmental trajectories in DIV70 forebrain organoids. **a**, Venn diagram depicting the overlap of EN lineage driver genes, DEGs identified in ENs (scEN DEGs) and target genes carrying SETD1A peaks. **b**, Latent time plots along velocity of a subset of high confidence representative examples of SETD1A target and non-target dysregulated EN lineage driver genes that display consistent changes in both bulk RNA-seq and scRNA-seq EN datasets in terms of both *P* value and fold-change direction. Five SETD1A target genes (NR2F1, NR2F2, VIM, NRP2 and LDHA) and three non-target genes (ACAT2, RBFOX1 and PCDH10) are depicted. **c**, Module enrichment across cell types (left) and their main functions of the genes included in each module (right). Nine major modules (M1-M9) were identified by WGCNA analysis and their sizes and colors were plotted based on –log_10_ (*P* value). **d**, Pie chart depicting portions of up- and down-regulated SETD1A target and non-target DEGs in different cell types along the EN lineage. Up_target denotes upregulated DEGs with SETD1A binding sites, Up_nontarget denotes DEGs without SETD1A binding sites. Total number of DEGs in each cell types are shown in brackets.

To obtain a more unbiased and comprehensive evaluation of the functional impact of the transcriptional dysregulation we also performed co-expression analysis on EN lineage cells from DIV70 WT organoids using the hdWGCNA pipeline. We identified 9 distinct Weighted Gene Correlation Network Analysis (WGCNA) modules involved in neuronal maturation in the EN lineage (**Fig. 5c**) (**Supplementary Table 8**). Each module contains multiple genes that map to a fraction of cell types in the EN lineage ((**Extended Data Fig. 12a**,b) and are enriched in specific GO terms and biological pathways ranging from chromatin remodeling to neuronal development and mitochondria-related functions (**Supplementary Table 8**). We calculated harmonized module eigengenes (a measure of the expression level and connectivity of the co-expression network) across cell groups within the EN lineage and queried for these gene modules in mutant and WT organoids to indicate if the module gene expression networks are up- or down-regulated. Each module is significantly altered in some but not all cell types (**Extended Data Fig. 12c**). In agreement with from our analysis, module M6 that contains multiple genes associated with neuronal development was consistently altered in mutant IPCs, nEN and EN cells (**Fig.5c**). Intriguingly, module M4, closely related to oxidative phosphorylation (OXPHOS) and cytoplasmic translation functions, showed the highest overlap with DEGs across IPC, nEN and EN with a sizeable portion of DEGs in these cell types contribute to this key co-expression module. Overlapping DEGs are for the most part upregulated in mutant cells and are enriched in functional terms relevant to “OXPHOS” and “aerobic respiration” (**Supplementary Table 8**). In terms of the number of modules and related functions disrupted, IPC-nEN cells displayed the greatest perturbation. Examination of DEG patterns and functional enrichment in each cell type in the EN lineage mirrored the finding of the WGCNA analysis and confirmed the pronounced perturbation of mutant IPC-nEN cells, which display the highest portion of upregulated SETD1A target DEGs and a pronounced disruption of gene networks involved in OXPHOS (**Fig. 5d and Supplementary Table 9**). Collectively, these findings indicate that disruption of *SETD1A* results in direct or indirect dysregulation of the temporal gene expression programs that establish and maintain specific cell states of the EN lineage.

### SETD1A mutation alters EN maturation trajectories and expression profiles

To determine if dysregulation of temporal gene expression programs and related alterations in EN development persist into later maturation, we performed scRNA-seq on organoids from mutant and WT iPSC lines at DIV150, during ongoing EN maturation. We obtained high-viability (>80%) single-cell suspensions from three organoids per line, and our quality control metrics confirmed uniform data quality (low mitochondrial and ribosomal gene counts; **Extended Data Fig. 13a-b**). As observed at DIV70, cell type quantification by Milo^42^ at DIV150 revealed a nominally significant lower proportion of differentiated ENs and a higher proportion of RGs in mutant organoids (**Fig. 6a**). Using Monocle3, we mapped EN lineage cells along a pseudotime trajectory defined by transcriptomic changes (**Fig. 6b** and **Extended Data Fig. 13c**). Compared to WT, mutant organoids exhibited a higher distribution of cells in early pseudotime (progenitor states) and a lower distribution in later pseudotime (mature states) (two-sided K-S test, p < 0.001; **Fig. 6c**).

**Figure 6.**
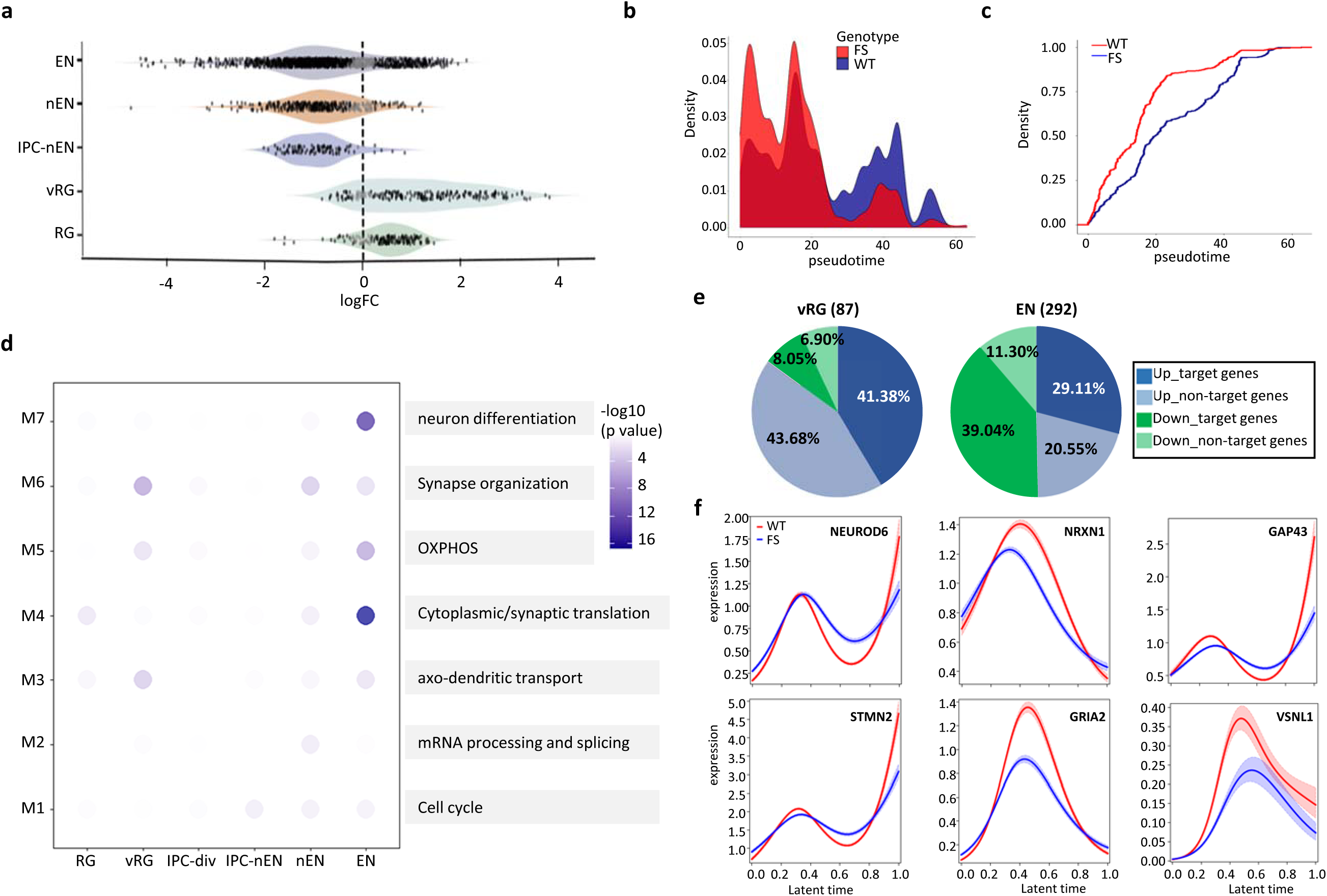
Aberrant gene expression and maturation trajectories in mutant forebrain organoids at DIV150. **a**, Beeswarm plots showing log fold change (Mutant/WT) in neighborhoods containing cells from the same cell type at DIV150. Mutant organoids have higher fraction of neural progenitor cells (RG and vRG) and lower fraction of differentiated ENs compared to WT organoids (spatialFDR <0.1) **b,** Pseudotime trajectory of the EN lineage as calculated with Monocle3. The ridge plot (left) shows that, at DIV150, mutant cells are predominantly distributed in the early, progenitor phase of the trajectory, whereas WT cell distribution is shifted towards the terminal phase of the trajectory represented by more mature ENs. **c,** K-S test (right) indicating a significant difference between the two distributions (P = 1×10^-5^). ECDF: Empirical Cumulative Distribution Function. **d,** Module enrichment across cell types (left) and the main functions of the genes included in each module (right). Seven major modules (M1-M7) were identified by WGCNA analysis and their sizes and colors were plotted based on –log10 (P value). **e,** Pie chart depicting the relative portions of up- and down-regulated SETD1A target and non-target DEGs in vRG and ENs. Up_target denotes upregulated DEGs with SETD1A binding sites, Up_nontarget denotes DEGs without SETD1A binding sites. Total number of DEGs in each cell types are shown in brackets. **f,** Latent time plots along velocity of a subset of representative examples of SETD1A target dysregulated DIV150 EN lineage driver genes that display dynamic temporal disparities between mutant and WT organoids.

We performed differential gene expression analysis across the EN lineage using the MAST package (**Supplementary Table 10)**. In ENs, the most abundant cell type in the DIV150 EN lineage, we identified 292 DEGs (145 up and 147 down, padj < 0.05, **Supplementary Table 10**) with 68% (199/292) being SETD1A targets. Among these, 19 contain a SETD1A DBS, 10 of which exhibit differential H3K4me3 signals and a strong positive correlation between differential gene expression and changes in SETD1A/H3K4me3 signals at the associated DBS (R² = 0.63 and 0.76; **Extended Data Fig. 13d**). Co-expression analysis on EN lineage from DIV150 WT organoids using WGCNA identified seven modules linked to neuronal maturation, each exhibiting differential dysregulation across subpopulations, with ENs showing the most pronounced perturbation in mutant organoids (**Fig. 6e, Extended Data Fig. 13e)**. Notably, 60% of EN DEGs overlapped with these modules. Module M6, associated with neuronal differentiation and synaptic organization, showed the highest overlap with DEGs in ENs, with a sizeable portion of EN DEGs (15%) contributing to this key co-expression module. Modules M5, related to OXPHOS and M4, related to protein translation followed closely, overlapping with 8.9% and 8.8% of EN DEGs, respectively (**Supplementary Table 11**). DEG patterns and functional enrichment in EN lineage cells (**Supplementary Table 10**) align with WGCNA findings, with downregulated genes in ENs primarily associated with neuronal generation and projection, while upregulated genes are enriched in mitochondrial electron transport and OXPHOS pathways. In progenitor vRG cells, the second most abundant cell type, DEGs were predominantly upregulated (74 up, 13 down), including those involved in OXPHOS. This transition from early upregulation of SETD1A target genes linked to metabolic disruptions to later, more balanced dysregulation affecting neurodevelopment parallels the expression dynamics observed in progenitor states of the DIV70 EN lineage (**Fig. 6d**).

To further explore the transcriptional alterations driving the aberrant maturation of mutant ENs, we leveraged Monocle3 pseudotime analysis and identified 78 putative lineage driver genes—those whose expression dynamics are likely to influence EN maturation from DIV70 to DIV150. Of these, 17 overlapped with DEGs in DIV150 ENs (**Supplementary Table 10**) and 14 of them were identified as SETD1A target genes. Several of these dysregulated SETD1A target driver genes are implicated in late neuronal maturation processes, including neuronal differentiation, synapse formation and transmission, axonal growth, dendritic arborization, and the establishment of neuronal connectivity. Notable examples include established markers such as NEUROD6, STMN2, GRIA2, and VSNL1, as well as NRXN1 and GAP43. The expression patterns of these target genes were markedly altered between WT and mutant organoids, with many displaying dynamic temporal disparities (**Fig. 6f**, **Extended Data Fig. 14a-b**). In summary, these results indicate that SETD1A LoF disrupts temporal gene expression programs coordinating key developmental and metabolic pathways, leading to aberrant neuronal maturation.

### Commonalities in EN lineage gene expression profiles

SETD1A mutations broadly impact gene networks and pathways in ENs, causing cell type- and developmental stage-specific changes. In this context, overlapping DEGs highlight key molecular processes consistently disrupted, providing insight into the core mechanisms affected during early development. Our analysis identified 49 genes that were consistently dysregulated in both DIV70 and DIV150 EN datasets, exhibiting uniform changes in fold-change direction (**Extended Data Fig. 14c**). Notably, 29 of these genes are direct SETD1A targets, with nine NGRN, PUS7L, GNPDA1, UBE2T, PTMA, NR2F1, VIM, MED30, MAP2) containing a SETD1A DBS, a significant enrichment compared to the overall SETD1A target gene set (31% versus 2.5%). Furthermore, five of these SETD1A DBS exhibit differential H3K4me3 signals (NGRN, PUS7L, GNPDA1, UBE2T, MED30) (**Supplementary Table 10**). These genes regulate diverse processes, such as neuronal development, structural integrity, metabolic functions, gene regulation and proteostasis. We highlight PUS7L and UBE2T as indicative of previously unrecognized pathways whose downregulation may contribute to the effects of *SETD1A* mutations on developmental trajectories. PUS7L is involved in RNA pseudouridylation, a modification that enhances RNA stability and function^49^, while UBE2T is crucial for protein homeostasis as well as DNA repair through its role in the Fanconi anemia pathway^50^. By comparison, only 13 genes showed consistent dysregulation in both DIV70 and DIV150 EN datasets, but with opposing directions. Of these, seven are direct SETD1A targets, with two (29%) containing a SETD1A DBS (NR2F2 and THSD7A; **Supplementary Table 10**).

In summary, our analysis of forebrain organoids has uncovered how *SETD1A* haploinsufficiency disrupts temporal gene expression patterns within the EN lineage, and highlighted distinct mechanisms by which SETD1A regulates gene expression at enhancers versus promoters. To assess the clinical relevance of these findings, we examined SETD1A binding profiles in the human fetal brain and explored the relationship between SETD1A-dependent genomic regulation and modalities linked to psychiatric disorders.

### SETD1A binding sites in EN of human fetal brain cortices

We isolated ENs from human fetal frontal cortices at 19–20 gestational weeks using a previously established FACS-based method^40^ (see Methods) and performed CUT&Tag for H3K4me3 and SETD1A (**Extended Data Fig. 15a**). A merged peak list from three biological replicates was generated and filtered to remove potential nonspecific peaks identified by the IgG negative control. We identified 15,387 H3K4me3 peaks and 13,211 SETD1A peaks in the fetal brain (**Fig. 7a, Supplementary Table 12**). The majority of H3K4me3 peaks (77.80%) were located in promoter regions, consistent with its well-established role as a promoter-associated histone mark. We observed an almost complete overlap of H3K4me3 peaks in ENs from fetal brain and forebrain organoids (**Extended Data Fig. 15b**), supporting the fidelity of H3K4me3 binding patterns in forebrain organoids.

**Figure 7.**
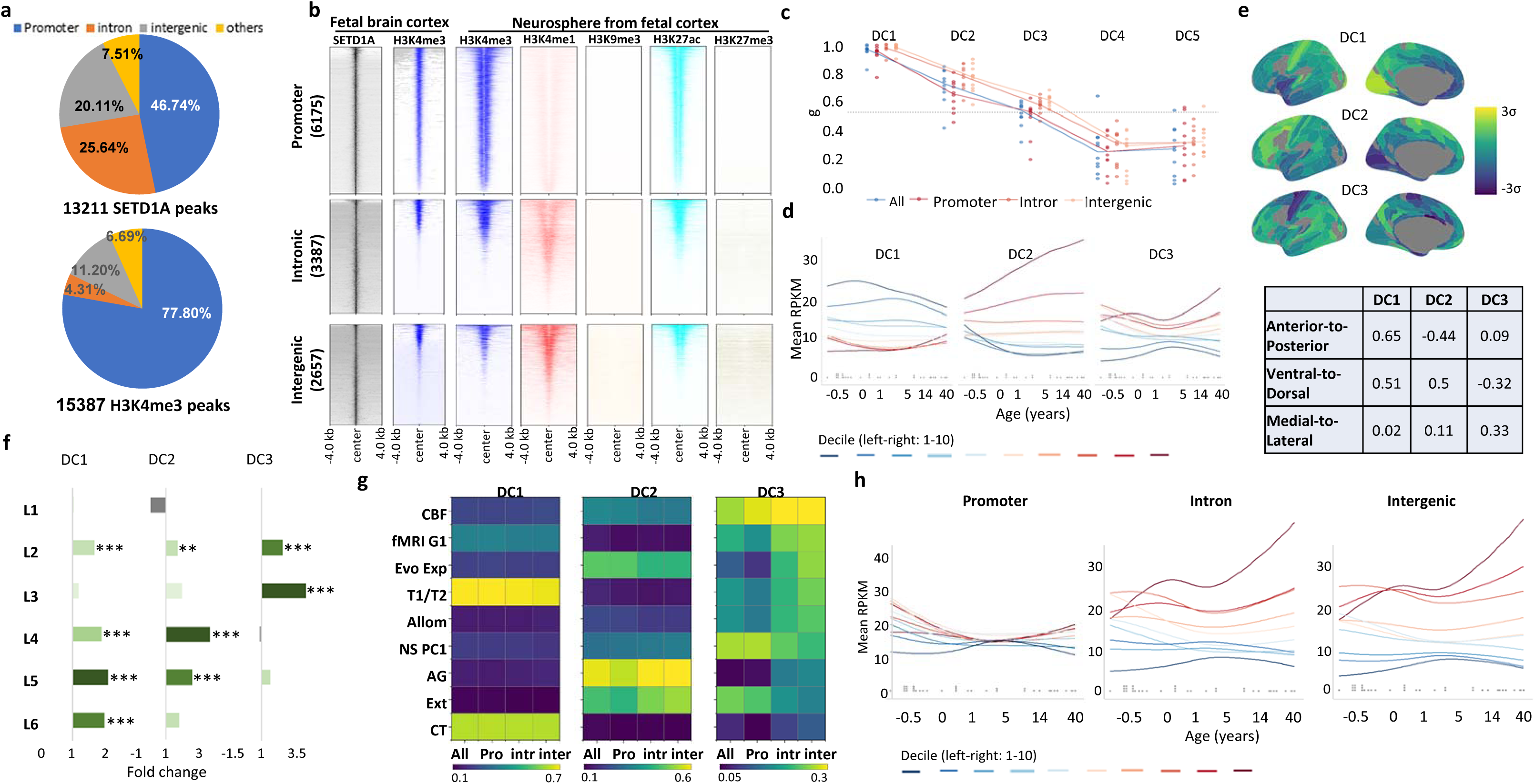
Identification and characterization of SETD1A target genes in EN from human fetal frontal cortices. **a**, Genomic distribution of SETD1A peaks (n = 13211) and H3K4me3 peaks (n = 15387) identified in excitatory neurons sorted from frontal cortices of healthy fetal brains at gestation week 19-20 (GW19-20). **b**, Heatmap of H3K4me3 mark identified in sorted excitatory neurons of human fetal forebrain at GW19-20, five histone marks (H3K4me1, H3K4me3, H3K9me3, H3K27ac and H3K27me3) centered on SETD1A peaks (n = 13211) across a ±2kb window. The five histone modification profiles were extracted from ENCODE datasets from human fetal (17 weeks) cortex-derived neurospheres and compared to the SETD1A genomic occupancy map. SETD1A peaks were clustered in promoter (<1kb) (top, 6175 peaks), intronic (middle, 3387 peaks) and intergenic regions (bottom, 2657 peaks). **c**, Identification of stable transcriptional components within SETD1A target gene lists (all targets, promoter targets, intronic targets, and intergenic targets) using DME. Generalizability (g) was assessed, with a threshold of g = 0.5 used for stable component selection. **d**, Developmental trajectories of all genes with SETD1A binding sites in DC1 to DC3 in the developing human brain. Gene expression levels were first fitted using regression for each gene, after which gene weights were averaged within each decile and represented by color-coded lines. Gray dots above the x-axis indicate the actual ages of the donor brains. RPKM refers to reads per kilobase per million mapped reads. **e**, DC1–DC3 components were orthogonally aligned in anatomical space based on Pearson’s correlations between regional scores and the XYZ coordinates of region centroids. The table presents the correlations of DC1– DC3 with each anatomical axis. DC1 and DC2 demonstrated alignment along the anterior-posterior axis in opposite directions. All three components (DC1, DC2, and DC3) exhibited alignment along the ventral-dorsal axis, with DC3 oriented in the opposite direction to DC1 and DC2. Notably, only DC3 showed alignment with the medial-lateral axis. **f**, Overlap between SETD1A target genes in each of three transcriptional components and cortical layer-specific marker gene-sets. **g**, Correlation heatmaps depicting the relationship between nine neuroimaging maps and the three transcriptional components (DC1–DC3) based on the AHBA dataset. “All,” “Pro,” “Intr,” and “Inter” represent all SETD1A target genes, as well as promoter, intronic, and intergenic subsets, respectively. **h**, Developmental expression profiles of genes with SETD1A binding sites in promoters, intronic regions, and intergenic regions within DC3. Gene expression levels were first regression fit for each gene and then gene weights were averaged within each decile and represented by color-coded lines. The gray dots above the x axis represent the actual ages of the donor brains. RPKM denotes reads per kilobase per million mapped reads.

In contrast, SETD1A peaks in ENs from fetal brain were equally distributed between promoter regions (46.74%) and enhancer-like regions, with 20.11% located in intergenic regions and 25.64% in intronic regions (**Fig. 7a**). SETD1A-bound promoters show significant H3K4me3 enrichment, aligning with its role in catalyzing H3K4 trimethylation at promoter regions. Further analysis of ENCODE histone modification profiles from 17-week human cortex-derived neurospheres, revealed that SETD1A binding sites are enriched for both promoter (H3K4me3, H3K27ac) and enhancer (H3K4me1, H3K27ac) marks, consistent with our organoid findings (p = 1 × 10⁻⁵, Z = 197.0 for H3K4me1; p = 1 × 10⁻⁵, Z = 446.4 for H3K4me3; p = 1 × 10⁻⁵, Z = 332.7 for H3K27ac by RegionR). In contrast, there was little to no enrichment for the repressive histone marks H3K9me3 (p = 1 × 10⁻⁵, Z = −11.0) and H3K27me3 (p = 1 × 10⁻⁵, Z = 9.8), based on data from the ENCODE repository (**Fig. 7b**).

We identified 5,201 SETD1A peaks overlapping between the fetal brain (n = 13,211) and forebrain organoids (n = 10,218), a significantly higher-than-expected overlap (p = 1 × 10⁻⁵, Z = 761.8, RegionR enrichment analysis). Overlapping peaks were primarily located in promoters (**Extended Data Fig. 15c**–d). Overall, there was a change in the distribution of SETD1A binding sites in the fetal brain compared to forebrain organoids, with a higher proportion localized to enhancer-like regions. This shift likely reflects differences in developmental stage and EN activity.

Two features of the genomic architecture of SETD1A binding identified in EN neurons from forebrain organoids were preserved in fetal brains. First, analysis of HiC-seq data from the cortical and subcortical plate of human fetal cerebral cortex^33^ found highly significant overlap between SETD1A binding sites and chromatin contacts represented by HiC loci (*P* = 2×10^-5^, Z = 10.3) (**Extended Data Fig. 15e**) further supporting the view that SETD1A may modulate 3D chromatin architecture. Second, *de novo* motif analysis using HOMER identified 24 significantly enriched TF motifs within SETD1A binding sites (FDR p < 10⁻^15^, **Supplementary Table 12**), collectively covering 96.7% (12771 of 13211) of the total SETD1A-bound regions. Notably, 6 of the top 10 enriched motifs overlapped with those identified in forebrain organoids, including RFX and SP4 motifs (**Extended Data Fig. 15f**) supporting the notion that SETD1A regulates cell type- and developmental stage-specific gene expression through interactions with a conserved set of auxiliary TFs in both the developing brain and forebrain organoids.

### SETD1A binding profiles influence both cellular processes and macroscale brain organization

We performed GO analysis to elucidate the distinct biological functions associated with SETD1A binding at various cis-regulatory regions in human fetal brains. Importantly, the GO term enrichment patterns observed were highly similar to those identified in forebrain organoids. Target genes with promoter-associated SETD1A binding sites showed the highest enrichment scores for terms related to “nuclear protein-containing complex” and “catalytic complex”. In contrast, target genes with intronic and intergenic SETD1A binding sites were enriched for terms related to “neuron projection” and “neuron development” (**Extended Data Fig. 15g**). Consistently, additional analysis of SETD1A target enrichment among markers of specific cortical layers revealed distinct patterns: genes with promoter-associated binding sites did not exhibit clear specificity to any particular layer, while genes with enhancer-associated sites were significantly enriched in marker genes of layers 2, 5, and 6 (**Extended Data Fig. 15h**).

To enhance the interpretability of our SETD1A target network in brain function and pathology, we leveraged human brain atlas datasets to identify core transcriptional components, assess their co-expression patterns within brain regional architecture, and investigate whether promoter-versus-enhancer distinctions within these components correlate with macroscale brain structure and function. To this end, we applied diffusion map embedding (DME) to the Allen Human Brain Atlas (AHBA) dataset, integrating it with comprehensive SETD1A target gene lists (all targets, promoter targets, intronic targets, and intergenic targets), which yielded three transcriptional components (DC1–DC3) with generalizability scores >0.5 (**Fig. 7c**, **Supplementary Table 13**). A total of 39.4% of SETD1A target genes were represented in the three components, with DC1 explaining 28.6% of the total variance of the dataset, DC2 explaining 9.4%, and DC3 explaining 7.2%, respectively.

The three components displayed modest yet statistically significant differences in the distribution of promoter, intronic, and intergenic regions, with DC3 exhibiting a slight elevation in intergenic enhancer-like regions (**Supplementary Table 13**). GO enrichment analyses of these components, stratified by positive and negative gene weights, revealed distinct functional associations. In DC1, negatively weighted genes were enriched for synapse-related processes and neuronal generation, while positively weighted genes were associated with transcriptional regulation and histone modification. In DC2, negatively weighted genes showed enrichment in DNA repair and dendritic functions, whereas positively weighted genes were linked to mitochondrial metabolic processes. In DC3, negatively weighted genes were associated with cytoskeletal dynamics, while positively weighted genes were enriched for synaptic signaling and glutamatergic synapse function (**Supplementary Table 13**). We further examined the developmental trajectory of these transcriptional components by analyzing RNA-seq data from the BrainSpan dataset, which includes samples from 35 donor brains spanning mid-gestation (8 post-conception weeks) to 40 years of age. This analysis revealed distinct temporal expression patterns: target genes within the DC1 component exhibited a predominantly prenatal bias, whereas those in the DC2 component showed a gradual increase in expression across development. In contrast, DC3 target genes displayed a primarily postnatal bias. Additionally, SETD1A target genes within these components demonstrated distinct axial alignments within anatomical space (**Fig. 7d-e**) as well as distinct layer-specific enrichment, with DC1 and DC2 significantly enriched in cortical layers 2, 4, 5, and 6, while DC3 target genes were predominantly enriched in supragranular layers 2 and 3 (**Fig. 7f**).

We examined the relationships between these transcriptional components and nine neuroimaging maps derived from MRI and PET data by aligning them with AHBA brain regions to investigate their contributions to structural, metabolic, and functional modalities along the sensorimotor-to-association axis of cortical organization^51^. While some modalities exhibited stronger correlations with a single component, others showed moderate associations with multiple components (**Fig. 7g**). For instance, markers of T1w/T2w myelination and cortical thickness were strongly associated with DC1 (r = 0.75, r = 0.68, respectively), while aerobic glycolysis, evolutionary cortical expansion, and externopyramidization were most strongly linked to DC2 (r = 0.62, r = 0.50, r = 0.48, respectively). In contrast, function-related modalities, including the principal gradient of functional MRI connectivity (fMRI G1), and first principal component of cognitive terms meta-analyzed by Neurosynth (Neurosyn P1) and cerebral blood flow (CBF) demonstrated similar modest correlations across all three components.

Notably, analysis of the promoter-enhancer distinction revealed that while DC1 and DC2 correlations were independent of SETD1A binding location, DC3 correlations varied by genomic context. Specifically, SETD1A target genes with intergenic and intronic binding sites showed stronger associations with certain neuroimaging modalities, whereas those with promoter binding sites exhibited weaker correlations. Although functional enrichment analysis of the DC3 component across promoter, intronic, and intergenic regions revealed broadly similar profiles for neuronal terms, each enriched in categories such as “neuron projection,” “synaptic signaling,” and “cell adhesion” (**Supplementary Table 13**) and all regions exhibited enrichment in supragranular layers, particularly in L3 for enhancer-bound targets (**Extended Data Fig. 15i**), we observed striking differences in their developmental expression profiles. Specifically, genes with promoter-bound SETD1A exhibit consistently low expression levels throughout development, whereas those with intronic and intergenic binding sites display robust postnatally-biased expression, particularly after adolescence (**Fig. 7h**). Overall, these findings underscore that the regulatory functions of SETD1A depend on both genomic context and developmental timing, influencing cellular processes as well as macroscale brain organization.

### SETD1A binding profiles are associated with disease-related neuroimaging deviations and genetic liability

To assess the clinical relevance of these transcriptional components in major neuropsychiatric disorders, we leveraged the BrainChart neuroimaging dataset, which includes over 125,000 MRI scans, to examine the relationship between these components—stratified by SETD1A binding profiles—and atypical regional cortical volumes in SCZ, ASD, and MDD. Notably, cortical shrinkage in SCZ was significantly associated with SETD1A intronic and intergenic target genes within the DC3 component, highlighting the potential role of enhancer-bound SETD1A targets in SCZ pathology (**Fig. 8a**). In contrast, no significant correlation was found between atypical regional cortical volumes in ASD or MDD and any of the transcriptional components.

**Figure 8.**
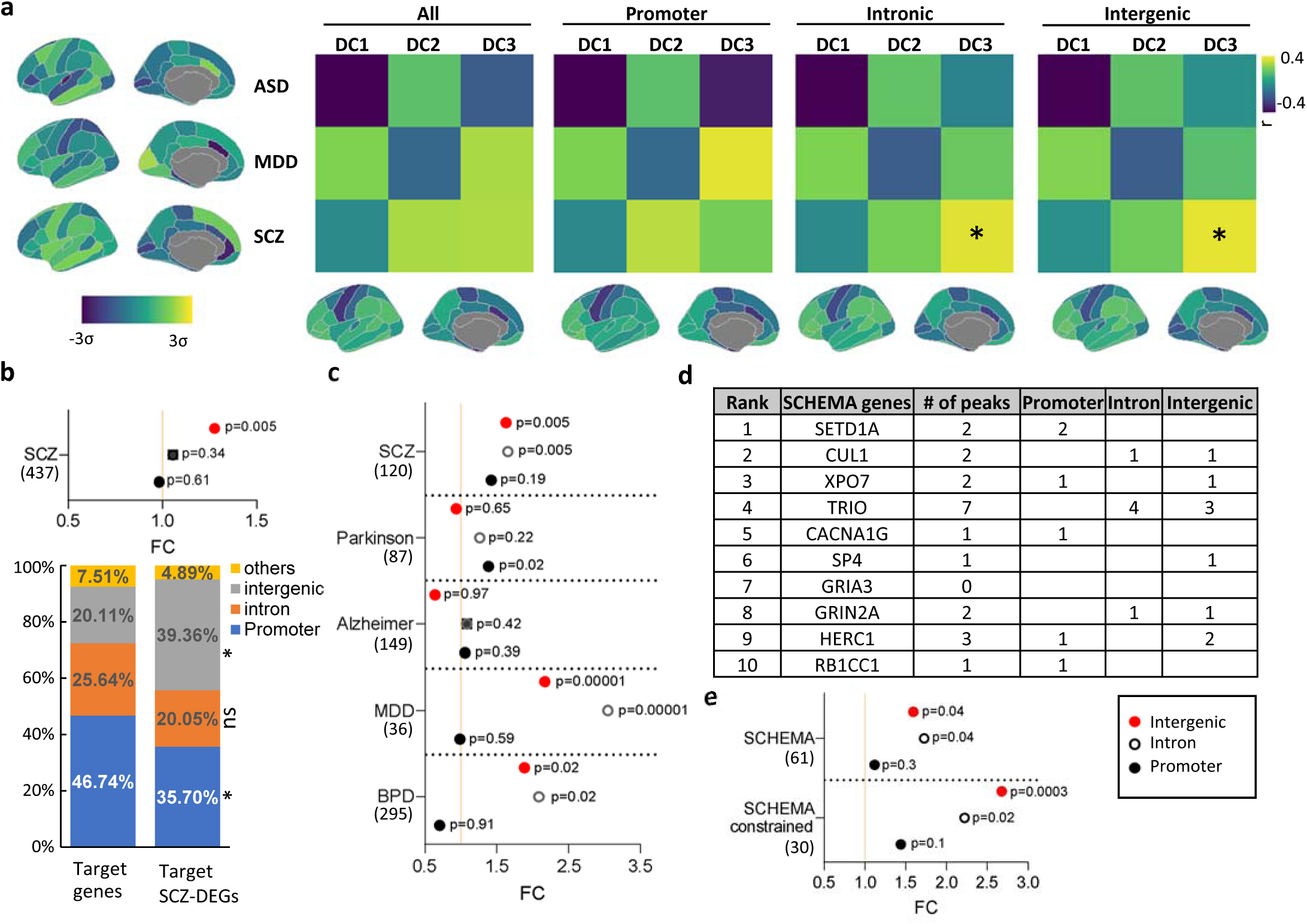
SETD1A binding profiles are associated with disease-related neuroimaging deviations and genetic liability. **a**, Spatial correlations between region-specific cortical volume changes associated with ASD, MDD and SCZ and the brain distribution of SETD1A targets with distinct genomic features (all targets, promoter targets, intronic targets, and intergenic targets, respectively) across the three transcriptional components. Yellow indicates positive correlation, while dark purple represents negative correlation. Statistical significance was assessed using two-sided FDR-adjusted spatially autocorrelated spin permutations, with corrections for multiple comparisons (* p < 0.05). Cortical volume shrinkage in ASD, MDD, and SCZ cases is depicted on the left, with yellow-highlighted brain regions indicating greater shrinkage, computed as z-scores derived from normative modeling of over 125,000 MRI scans. The cortical maps at the bottom illustrate the brain-wide distribution of SETD1A target genes with distinct genomic features within the DC3 component, projected onto the Desikan-Killiany parcellation. **b**, Forest plot illustrating the enrichment of SETD1A target genes with distinct genomic features within a consensus DEG list derived from independent SCZ studies (top). Permutation analysis (bottom) reveals a significant shift in the genomic distribution of intergenic and promoter regions among SCZ DEGs compared to the overall distribution of SETD1A binding sites in the fetal brain. **c**, Forest plot illustrating the enrichment of SETD1A target genes with distinct genomic features within prioritized genes mapped to common variants associated with SCZ, MDD, BPD, Parkinson’s disease and Alzheimer’s disease. **d**, Forest plot illustrating the enrichment of SETD1A target genes with distinct genomic features within SCHEMA genes (n = 61, FDR < 0.001) and constrained SCHEMA genes (n = 30, pLI>0.9).

Follow-up comparisons with gene expression and human genetic data yielded results consistent with our neuroimaging findings. Specifically, we first compared SETD1A target genes with a consensus DEG list derived from independent RNA-seq studies of dorsolateral prefrontal cortex tissue in SCZ. We found that SCZ DEGs (n = 241) are significantly enriched in genes with SETD1A-bound enhancer-like regions (**Fig. 8b, upper**) suggesting that variations in the SETD1A levels may contribute to disease-associated transcriptional changes even in individuals without SETD1A mutations. Accordingly, SETD1A peaks overlapping SCZ DEGs are more frequently located in intergenic regions (39.96% versus 20.11%, p <0.05), and less often in promoters (35.70% versus 46.74%, p <0.05) compared to overall SETD1A binding sites (**Fig. 8b, lower**).

Furthermore, using data from a recent SCZ GWAS^52^, we found that SETD1A peaks overlapping with risk genes mapped to common variants significantly associated with SCZ were enriched in intergenic and intronic but not in promoter regions. Among the 120 genes prioritized by SCZ-GWAS, 40 genes contained one or more SETD1A binding sites in intergenic and intronic regions. These genes are enriched in functional terms pertinent to brain development, synaptic structure, and function (**Supplementary Table 14**). Control analyses showed that genes mapped to common variants associated with Alzheimer’s and Parkinson’s disease were not enriched in any genomic compartment, suggesting a more specific role for SETD1A in the pathogenesis of psychiatric disorders (**Fig. 8c**). Interestingly, a pattern of enrichment similar to that observed in SCZ was also found for genes mapped to common variants associated with major depressive disorder (MDD, n = 295 genes) and BPD (n = 36 genes), consistent with GWAS studies identifying a shared genetic aetiology between MDD, BPD and SCZ^53,54^ as well as recent suggestive evidence implicating SETD1A in genetic risk for BPD^55^.

Expanding our enrichment analysis, we investigated whether SETD1A target genes are overrepresented among genes implicated in SCZ through rare-variant burden analyses. Utilizing data from the SCHEMA consortium’s case-control study of 24,248 SCZ patients and 97,322 controls, we found that among the 10 genome-wide significant SCZ risk genes (p < 2.2 × 10⁻⁶; odds ratios: 3–50), nine contained at least one SETD1A binding site (**Fig 8d**, **Supplementary Table 14**). Notably, TRIO exhibited the highest number of SETD1A binding sites (n = 7) with four in intronic regions and three in intergenic regions. Extending this analysis to genes identified at varying FDR thresholds (<0.05, 0.01, 0.005, and 0.001), we observed a positive correlation in the overlap between genes in each group and SETD1A targets in intergenic and intronic regions, but not in promoter regions. We investigated the role of gene constraint, i.e., the intolerance to mutational changes for genes frequently affected across neuropsychiatric disorders^56^ and found that this correlation was even more pronounced when focusing on constrained genes (pLI ≥ 0.9, **Extended Data Fig. 16**). Consistently, SETD1A targets overlapping SCHEMA genes identified at an FDR < 0.001 (n = 61), were significantly enriched in intergenic and intronic, but not in promoter binding (**Fig. 8e**). We identified 21 target SCHEMA genes harboring SETD1A enhancer binding sites (**Supplementary Table 14)** including, in addition to the genome-wide significant SCZ risk genes, genes encoding cell adhesion molecules, such as NEGR1; ion channels, such as KCNN3 and CACNA2D1; protein kinases, such as FYN; genes involved in DNA damage repair, such as SLF2; and genes associated with epigenetic regulation, such as ZMYND11 and KDM6B. Notably, we also identified AKAP11, a gene linked to BPD through exome sequencing studies^57^.

The minimal overlap between the SCZ DEGs, GWAS and SCHEMA risk genes underscores the widespread role of SETD1A-bound enhancer regions in modulating gene networks contributing to the complex genetic architecture of SCZ. Notably, minimal overlap was observed between SETD1A target disease genes and DEGs in the DIV70 and DIV150 EN lineages (7 and 3 genes, respectively, for the GWAS list and none for the SCHEMA list, **Supplementary Table 14**)—suggesting that most SETD1A target risk genes are not transcriptionally altered during early brain development, although scRNA-seq may have limitations in capturing all DEGs.

## Discussion

In this study, we generated forebrain organoids from iPSCs carrying a *SETD1A* LoF risk mutation to identify SETD1A genomic targets, characterize their properties and use this information to provide a molecular read-out of how *SETD1A* disease risk mutations affect the expression of ensemble of genes important for development of human cortical ENs. Chromatin profiling via CUT&Tag, combined with RNA sequencing, revealed numerous SETD1A binding sites across human regulatory elements, enabling the identification of high-confidence target genes as well as potential SETD1A recruiters and interactors that mediate the effects of SETD1A on early human neurodevelopmental processes. Analysis of these targets and transcriptomic changes induced by a SETD1A SCZ risk mutation uncovered robust molecular signatures indicative of disruptions in the proper temporal execution of neuronal differentiation and maturation programs in the EN lineage. Experimental validation, in two independent clonal lines, confirmed alterations in cell abundance, neurogenesis, differentiation, and neuronal excitability between mutant and WT organoids, findings predicted by our genomic and transcriptomic analyses, reinforcing the robustness of our results. These convergent molecular and cellular findings, consistently observed across organoid batches, rule out random developmental variability and highlight specific EN gene ensembles and biological pathways—including key regulatory and metabolic processes^48,58^—that are critically affected by SETD1A mutations.

Our assessment of the genomic binding profile of SETD1A revealed that reduced SETD1A levels in mutant neurons do not alter its global binding distribution but induce local changes at a subset of SETD1A-bound peaks, resulting in both diminished and enhanced binding. Comparison with corresponding changes in H3K4me3 levels showed a strong positive correlation between SETD1A occupancy and H3K4me3 enrichment. Notably, among the differentially bound peaks, a consistent pattern emerged across multiple DEG datasets, indicating that shifts in SETD1A occupancy are positively correlated with changes in both H3K4me3 levels and gene expression direction. These findings suggest that, for a subset of SETD1A binding sites, the mutation induces transcriptional changes by modulating the H3K4me3 epigenetic mark. While DBS constitute approximately 2.5% of the overall SETD1A target gene set, their representation increases to around 14% among SETD1A target DEGs in sorted DIV70 EN and is even higher when considering high-confidence target DEGs shared across all bulk and scRNA-seq EN datasets from DIV70 and DIV150. Importantly, differentially bound peaks are substantially enriched among genes whose expression dynamics likely govern EN development and maturation, especially those exhibiting pronounced temporal expression changes, suggesting that these alterations reflect a direct effect of altered SETD1A binding rather than secondary transcriptional consequences. Although our CUT&Tag sequencing may underestimate the total number of DBS due to sensitivity limitations, these findings indicate that a subset of SETD1A-bound sites exhibit differential sensitivity to LoF mutations, thereby identifying a set of high-confidence target genes directly impacted by these mutations.

Analysis of SETD1A binding sites in EN of human forebrain organoids and fetal brain cortices revealed important differences in the mechanisms by which SETD1A engages enhancer versus promoter elements to regulate target gene expression. Specifically, compared to SETD1A binding overall, peaks overlapping differentially expressed genes in mutant neurons are more frequently localized to enhancer regions. Additional analysis found that in contrast to predominately promoter binding in the overall SETD1A occupancy map, genomic sites with altered SETD1A occupancy in mutant neurons are highly enriched in intronic and intergenic enhancer elements. These shifts suggest that genes with SETD1A-bound enhancers may be relatively more vulnerable to the reduction of SETD1A levels due to risk LoF mutations than those regulated via promoters. This is possibly because of differences in the stability of the complexes that SETD1A forms in enhancers versus promoters influenced region-specific protein-protein interactions or local chromatin context^59^. Notably, target genes with promoter-bound SETD1A are enriched for basic cellular functions without cortical layer specificity, whereas those with binding sites in intergenic and intronic regions are predominantly enriched for neuronal functions and exhibit cortical layer specificity. Moreover, SETD1A’s regulatory effects correlate distinctly with neuroimaging maps and macroscale brain organization, depending on its genomic binding location. These distinctions are critical for elucidating how SETD1A mutations elevate neuropsychiatric risk and for developing targeted treatments with fewer side effects by specifically modulating SETD1A action at enhancer elements^14^.

Indeed, by triangulating evidence across datasets of cortical transcriptional architecture, neuroimaging, DEGs, GWAS and exome sequencing, we demonstrate that atypical phenotypes and the genetic risk of SCZ are predominantly associated with SETD1A enhancer targets. Notably, cortical shrinkage in SCZ is significantly associated with a specific component of SETD1A enhancer target genes that exhibit a postnatal expression bias, predominantly in neurons of the supragranular cortical layers, which are distinguished by dense cortico-cortical connections, late stage maturation, disproportionate expansion in humans compared to other species, and have been implicated in SCZ pathophysiology^60–62^.SETD1A enhancer-bound genes also show a highly significant overlap with established SCZ DEGs and risk gene sets. These findings suggest that reduced SETD1A levels in mutation carriers, or variation in its expression in sporadic cases, may lead to clinical deficits through a precisely orchestrated spatiotemporal pattern of gene expression that encompasses numerous disease-related genes.

In the context of SETD1A mutations, previous studies have begun to elucidate the neurodevelopmental consequences in mutant neuronal cultures^14–16,36,38,63,64^ and documented various phenotypic alterations in iPSC-derived neurons harboring *SETD1A* LoF mutations including altered dendritic complexity, synaptic transmission and plasticity as well as neuronal excitability. However, our study is the first to detail the developmental trajectory of iPSC-derived mutant neurons under biologically relevant conditions. By leveraging a forebrain organoid culture system that supports spontaneous differentiation into diverse cell types, we provide a more representative model of human neurodevelopment and link these processes to disrupted genomic and transcriptomic profiles. Our findings are largely consistent with previous reports, yet they uniquely reveal impaired EN neurogenesis and maturation, as well as recapitulate the effects of SETD1A haploinsufficiency on neuronal excitability.

In the context of SCZ, the primary psychiatric manifestation in adults with *SETD1A* mutations, our findings align with reports of impaired NPC development, from studies in patient neuronal models^65^. Although many such studies face challenges, including genetic heterogeneity and difficulty in precisely identifying neuronal subtypes, existing evidence suggests that NPCs struggle to differentiate into mature neurons. This may indicate that either a reduced initial NPC pool or impaired differentiation contributes to decreased neural density later in the disease, with underlying molecular causes varying among patients due to the genetic heterogeneity of SCZ^66,67^.

Our findings add to the discourse on the developmental origins of psychiatric disorders and may extend to other genetic causes of SCZ, particularly those with large-effect mutations that disrupt early brain developmental trajectories. It is conceivable that early developmental perturbation of the EN lineage emerging due to the *SETD1A* mutations, irrespective of whether it is transient or persisting throughout prenatal and early postnatal development, leads to changes in developing brain connectivity and activity patterns^68,69^, which may irreversibly propagate to later stages with long-lasting effects, and finally contribute to the psychiatric and cognitive symptoms in adulthood. However, our analysis suggests that additional mutational impact emerging at later developmental stages, affecting molecular and neural processes in EN in the mature brain through SETD1A target genes not expressed or transcriptionally altered during early development, also contribute to the risk conferred by SETD1A mutations. Our comprehensive identification of high-confidence SETD1A genomic targets, their properties and their overlap with established lists of disease-related genes and neuroimaging maps can serve as a foundation for further mechanistic and translational research aiming to elucidate the role of SETD1A in disease mechanisms and design targeted treatments.

## Supporting information

SETD1A and H3K4me3 Peaks Identified in Sorted Cortical Neurons from WT and Mutant Forebrain Organoids at DIV70

Publicly Available Histone Modification Profiles, GO Analysis and Human-Gained Enhancer Interaction Profiles in Fetal Cerebral Cortices

Motif Analysis of SETD1A Peaks and Functional Categorization of Target Genes

De Novo Motif Analysis and Genomic Annotation of SETD1A Peaks

Differentially Expressed Genes in Mutant Neurons, SETD1A Peak Overlap, and Enriched Motifs

Differential Binding Strength of SETD1A and H3K4me3 Peaks in Sorted Cortical EN and Their Overlap with DEGs

Differential Gene Expression and Driver Gene Identification in Distinct Cell Types of Forebrain Organoids at DIV70

Expression Profiles, GO Analysis, Distribution Comparison of WGCNA Modules in EN Lineage, and Overlap Between DEGs and Module Genes

GO Analysis of DEGs in IPC_nEN, nEN, and ENs at DIV70 and Upregulated Genes Enriched in OXPHOS in IPC_nEN

DEGs and Driver Genes in EN at DIV150 and Overlap Analysis Between DIV150 and DIV70 DEGs

Expression Profiles, GO Analysis, and Overlap of DEGs with WGCNA Modules in DIV150 EN lineage

Analysis of SETD1A and H3K4me3 Peaks in Human Fetal Brain, Genomic Annotations

Gene Lists and GO Enrichment Analysis of SETD1A Target Components (DC1-DC3) and Correlations with Neuroimaging Maps

Cortical Layer Specificity of SETD1A Target Genes within DC1-3 and Overlap with SCZ-Associated Genes

## Acknowledgements

This work was supported by the Stavros Niarchos Foundation (SNF), National Institute of Mental Health grant 5R01MH112860 and National Institute of Health-NCATS grant (UG3/UH3TR002151). Z.S was partially supported by a 2019 NYSTEM training grant and a 2022 NARSAD Young Investigator Grant from the Brain and Behavior Research Foundation. We thank the members of the Zhang laboratory for their technical assistance and providing pA-Tn5 for CUT&Tag assays. We thank Columbia Center for Translational Immunology (CCTI) for assistance with cell sorting, and the Columbia Stem Cell Initiative (CSCI) Flow Cytometry Core for assistance with flow cytometry. We thank Columbia University Genome Center for support with DNA and RNA library sequencing.

## Author contributions

Z.S, B.X and J.A.G designed the experiments and interpreted results from experimental assays. B.X and J.A.G supervised the work with contributions from S.M and S.K. Z.S performed all cellular, sequencing and chromatin assays. Z.S, J.F and B.X analyzed all sequencing and chromatin assays. H.Z and Y.S cultured hiPSCs and forebrain organoids. B.L performed whole-cell electrophysiological recordings. X.H and Z.W contributed to the delineation of transcriptional components and their correlation with macroscale brain structure and function. Z.Z contributed to the design and interpretation of the CUT&Tag assays. Z.S., B.X., J.A.G prepared the manuscript with contributions from all co-authors.

## Methods

### hiPSC line generation and characterization

For the generation of the FS mutation (c.1272delC), guide RNAs were designed using the CRISPOR tool (http://crispor.tefor.net/crispor.py). Efficacy of multiple guide RNAs was first evaluated in HEK293T and one guide RNA (5’-AGGCAGGAGGCCGGTAGTCC-3’) was selected. Ribonucleoprotein (RNP) complexes were made by mixing 100pmol HiFi-Cas9 with crRNA and tracrRNA (Integrated DNA Technologies, Inc) at a molar ratio of 1:3:3 at room temperature for 10 min. hiPSCs (MH0159020) from the NIMH stem cell repository with well-established stem cell properties were cultured in mTeSR™1, electroporated with RNP complex using a NucleofectorTM 2b Device (Lonza nucleofector) and then plated to 4 wells of 24-well plate. Approximately one week after electroporation, a sequential enrichment strategy was applied: A portion of cells in these four wells was subjected to Sanger sequencing for genotyping analysis individually, the resultant chromatograms were analyzed by Synthego (https://www.synthego.com/) for quantifying the percent of ACT/AT deletion. Cells with >2% ACT/AT deletion continued to grow until colonies formed, these colonies then subjected to further genotyping analysis as described above. Positive colonies with >15% ACT/AT deletion were subjected to single cell cloning by plating cells on 96-well plates at 1 cell/well density. Two weeks later, WT and mutant colonies were selected for subsequent analysis after confirming their genotypes. All hiPSC lines were maintained at low passage, they were karyotypically normal (as determined by G-banded karyotyping performed at Columbia Stem Cell Core Facility) and retained their stemness as confirmed by immunostaining for stem cell markers NANOG and SSEA-4.

### Antibodies

The following antibodies were used: *<u>Western blots:</u>* hSETD1A (rabbit, 1:1000, 61702S, Cell Signalling), hSETD1A (mouse, 1:600, SC-515590, Santa Cruz Biotechnology, Inc.), Histone H3 (rabbit, 1:30000, ab1791, abcam), Lamin B1 (rabbit, 1:1000, PA1048, Boster Bio), GAPDH (mouse, 1:1000, ab8245, abcam), goat anti-rabbit IgG (H+L) secondary antibody (1:5000, 65-6120, Thermofisher), goat anti-mouse IgG (H+L) secondary antibody (1:10000, A8924, Sigma).*<u>Immunocytochemistry and immunohistochemistry:</u>* The following antibodies were used: NeuN (mouse, 1:100, MAB377, Millipore), NeuN (rabbit, 1:500, ABN78, Millipore), SSEA4 (mouse, 1:500, ab16287, Abcam), NANOG (rabbit, 1:500, 4903, Cell signalling), DAPI (1:1000, D1306, Invitrogen). All secondary antibodies (goat, Molecular Probes) were used at 1:500 dilution for immunocytochemistry and immunohistochemistry, respectively. *<u>CUT&Tag analysis:</u>* IgG (rabbit, ab37415, abcam), hSETD1A (rabbit, ab70378, Abcam)

### Co-immunoprecipitation

Nuclear extract was prepared from HEK293Tcells using the Nuclear Complex Co-IP Kit (Active Motif. Inc.). For co-immunoprecipitation analysis, 500 μg of nuclear extract were incubated with 10 μg of rabbit anti-SETD1A antibody (ab70378, Abcam) or 10 μg of normal rabbit IgG (ab37415, abcam) at 4°C overnight. Then lysates were incubated with dynabeads protein A (10002D, Thermo Fisher) at 4 °C for 1hr. For immunoblot analysis, we used anti-hSETD1A (mouse, 1:600, SC-515590, Santa Cruz Biotechnology, Inc.) as the primary antibody and goat anti-mouse IgG (HRP) (1:10000, A8924, Sigma) as a secondary antibody.

### Immunocytochemistry

hiPSC were fixed with 4% paraformaldehyde (PFA) in phosphate-buffered saline (PBS) (pH 7.4) for 15 min at room temperature (RT), washed 3 times for 5 min in 1X PBS, permeabilized in 0.2% Triton X-100 in PBS for 10 min and washed again in PBS (3X for 5 min each time). Coverslips were blocked for 1hr with 10% horse serum (26050070, Gibco) PBS and incubated for 1 hr with primary antibodies diluted into blocking buffer (2% horse serum in PBS) and washed in PBS (3X for 5 min each time). Secondary antibodies were applied in blocking buffer for 1hr at RT. Following 3 times wash in PBS, the coverslips were mounted onto glass slides (47100, Richard-Allan Scientific).

### Forebrain organoid culture, viral labeling and FACS assays

Generation and maintenance of forebrain organoids was performed as described previously^70^. In brief, hiPSCs were dissociated with 0.5 mM EDTA in the incubator and triturated with 1ml tips into single-cell suspension. Single cells were then seeded into ultra-low-attachment 96-well plate (Nunc) at a density of 1 x 10^4^ cells per microwell, to form embryoid bodies (EBs) in medium containing mTeSR™1, 1 µg/ml heparin and Penicillin-Streptomycin antibiotics. EBs were cultured in medium containing ROCK inhibitor Y27632 (50 µM) for the first 24 h, followed by 5 days without interference in 96-well plates. On day 5, the medium was switched to medium containing DMEM/F12 (1:1) (Gibco; 11330), Glutamax (Invitrogen 35050-061), MEM-NEAA (Invitrogen), N-2 supplement (Gibco; 17502-048), Penicillin-Streptomycin, 0.1 mM β-Mercaptoethanol, Dorsomorphin (2 µM), SB431542 (10 µM) and IWR1e (3 µM). On days 25-39, 0.2% Chemically Defined Lipid Concentrate (Gibco; 11905031) was added to the medium. Medium was changed twice per week. On day 40, EBs were transferred to ultra-low attachment 24-well plates. On day 40-80, EBs were maintained in medium containing DMEM/F12 (1:1) [supplemented with Glutamax, MEM-NEAA, N-2 supplement, Penicillin–Streptomycin, 0.1 mM ®-Mercaptoethanol, 10% fetal bovine serum (FBS), and 1% Matrigel (Corning)] with weekly medium change. After day 80, Matrigel was added bi-weekly to the medium.

Forebrain organoids were transduced with the AAV-hSYN-GFP virus (Addgene, 105539-AAV1) at DIV55, to label neurons with GFP. Sorting of neurons labelled with GFP at DIV70 was conducted using a BD influx cell sorter in 1.5 ml tubes pre-coated with 10% FBS. Sorted neurons were then collected by centrifugation at 400g at 4°C for 10 min, followed by ice-cold PBS wash. Cell pellets were processed for CUT&Tag and RNA-seq simultaneously.

### Flow cytometry

Forebrain organoids at different developmental stages were first dissociated into single cell suspensions using papain digestion (Worthington Biochemical), followed by fixation and permeabilization with eBioscience™ Foxp3/Transcription Factor Staining Buffer Set (Thermofisher). Primary antibody (Ki67, SOX2, CTIP2) incubation was conducted at 4°C overnight. Following a washing step with PBS, cells were incubated with 1:500 Alexa Fluor secondary antibodies (Thermofisher) for 1hr at room temperature. Analysis was performed on a NovoCyte (Penteon) flow cytometer. 100-500K events were acquired for each sample with fluorescence measured in logarithmic scale. Cells incubated only with secondary antibodies served as background fluorescence, and were used to set the gating parameters. After exclusion of cell aggregates and small debris by forward and side scatter gating, data were analyzed using the NovoCyte software and plotted in a histogram format. Fluorescence gates were set below 2% of blank histogram and events corresponding to a fluorescence signal exceeding this percentage were considered as positive events.

### Whole-cell electrophysiological recordings

hiPSC lines were first cultured in mTeSR™1 on Matrigel-coated plates, and then switched to Stemflex medium (STEMCELL Technologies). hiPSCs were differentiated according to the previous protocol with some modifications^71^. Briefly, approximately 8000 hiPSCs were plated in an ultra-low attachment 96-well V-bottom plate (S-Bio). Two days later, cells were switched to neural induction medium (Advanced DMEM/F12 supplemented with 1% N-2 supplement, 1% penicillin/streptomycin and 2 µg/ml heparin) with the medium change every two days. A week later, embryoid bodies (EBs) were transferred to 6-well plates coated with 20 µg/ml laminin (Sigma), and medium was changed twice. At DIV14. EBs were passaged in NPC medium (Advanced DMEM/F12 supplemented with 1% N-2 supplement, 2% B-27 supplement, 1 µg/ml laminin, 20 ng/ml bFGF and 1% penicillin/streptomycin) with medium change 3 times per week, and EBs were passaged once per week. Fluorescence-activated cell sorting (FACS) was performed at passage 3 to obtain a pure population of CD24^+^/CD184^+^/CD44^-^/CD271^-^ NPCs with the protocol adopted from Yuan and colleagues^72^. From passage 5 onwards, NPCs were used for neural differentiation on 18 mm coverslips coated with poly-L-ornithine and subsequently with drops of 50 µg/ml laminin. NPCs were seeded in 50 microliter drops in neural differentiation medium (Neurobasal medium supplemented with 1% N-2 supplement, 2% B-27 supplement, 1 µg/ml laminin, 20 ng/ml BDNF, 20 ng/ml GDNF, 200 µM Ascorbic Acid, 1 µM cAMP and 1% penicillin/streptomycin) with medium change 3 times per week. Seeding density was optimized to achieve similar density. After 4 weeks, only half of the medium was replaced for medium change. Whole-cell patch-clamp recordings were performed after 8 weeks of NPC differentiation as previously described^73^. Briefly, cultures were equilibrated to oxygenated artificial cerebrospinal fluid (ACSF, 110 mM NaCl, 2.5 mM KCl, 2 mM CaCl_2_, 10 mM glucose and 1 mM NaH_2_PO_4_, 25 mM NaHCO_3_, 0.2 mM ascorbic acid and 2 mM MgCl_2_, pH 7.4). In the recording chamber, slides were continuously perfused with ACSF at 1.5–2 ml/min, saturated with 95% O_2_/5% CO_2_ and maintained at 20 – 22°C. Recordings were performed with borosilicate glass recording micropipettes (3–6 MΩ). Data were acquired at 10 kHz using an Axon MultiClamp 700B amplifier (Molecular Devices), filtered at 3 kHz, and analyzed using pClamp 10.1 (Molecular Devices). Current-clamp recordings were performed at a holding potential of −70 mV. Intrinsic membrane properties were analyzed using a series of hyperpolarizing and depolarizing square wave currents (500 ms duration, 1 s interstimulus interval) in 5 pA steps, ranging from −30 to +30 pA. Data analysis was performed using a custom-designed script in Igor Pro-8.0 (WaveMetrics). Input resistance was calculated from the first two hyperpolarizing steps. Active properties were extracted from the first depolarizing step resulting in AP firing. AP threshold was defined by the moment at which the second derivative of the voltage exceeded the baseline. AP amplitude was measured from threshold. Neurons were categorized as “firing” if they were capable of firing three or more mature APs without significant accommodation during a depolarizing current step. Voltage-clamp recordings were performed at a holding potential of −80 mV.

### Bulk RNA sequencing

Total RNA was extracted from sorted GFP positive neurons at DIV70 using RNeasy Mini Kit (Qiagen). Quality of RNA samples was assessed using a Bioanalyzer (Agilent 2100 bioanalyzer) to ensure the RNA Integrity Number (RIN)>9.0 for all samples. RNA-seq libraries were prepared from ∼400 ng of total RNA using the STRYPOLYA library prep kit (Illumina) and were sequenced at the Columbia Genome Center on a Novaseq 6000 instrument (Illumina) with 100bp paired-end reads. We used RTA (Illumina) for base calling and bcl2fastq (Illumina, version 2.17) for converting BCL to fastq format, coupled with adaptor trimming. We mapped the reads to the reference human genome (UCSC/hg38) using STAR ((Dobin et al., 2013) (version 2.5.4b). Read counts and normalization of the read count data and analysis of DEGs was performed using DESeq2^74^ (version 1.10.2) with default parameters.

### CUT&Tag assays

CUT&Tag libraries were prepared as previously described^27^ with some modifications. Briefly, 350K sorted neurons from forebrain organoids were harvested and gently washed once in wash Buffer (20 mM HEPES pH 7.5; 150 mM NaCl; 0.5 mM Spermidine; 1× Protease inhibitor cocktail) then resuspended in 100 µL Dig-wash Buffer (20 mM HEPES pH 7.5; 150 mM NaCl; 0.5 mM Spermidine; 1× Protease inhibitor cocktail; 0.01% Digitonin) containing 2 mM EDTA and a 1:100 dilution of the appropriate primary antibody. Primary antibody incubation was performed on a nutator platform overnight at 4 °C. Cells were washed 3 times with Dig-wash Buffer. An appropriate secondary antibody (Donkey anti-Rabbit IgG antibody for a rabbit primary antibody) was diluted 1:100 in 100 µL Dig-Wash buffer and cells were incubated at RT for 1 hour. Following 3 washes, cells were incubated with pA-Tn5 adapter complex (1:200 dilution) in 50 µL Dig-300 Buffer (0.05% Digitonin, 20 mM HEPES, pH 7.5, 300 mM NaCl, 0.5 mM Spermidine, 1× Protease inhibitor cocktail) at RT for 1 hour. Following another 3 washes, cells were resuspended in 200 µL Tagmentation buffer (10 mM MgCl2 in Dig-med Buffer), incubated at 37 °C for 1 h and then subjected to the addition of 2.25 µL of 0.5 M EDTA, 2.75 µL of 10% SDS and 0.5 µL of 20 mg/mL Proteinase K to stop tagmentation. Cells were then incubated at 55 °C for 2 hours followed by 85 °C for 20 min to inactivate Proteinase K. Genomic DNA was extracted using the ChIP DNA Clean & Concentrator kit and subjected to PCR amplification for library construction with barcoded i5 and i7 primers, using 25 µL NEBNext HiFi 2× PCR Master mix (NEB) and the following cycling conditions: 72 °C for 5 min (gap filling); 98 °C for 30 s; 12-14 cycles of 98 °C for 10 s and 63 °C for 30 s; final extension at 72 °C for 5 min and hold at 8 °C. Libraries were cleaned up using Ampure XP beads (Beckman Counter), then sequenced on Illumina NextSeq 550 sequencer with pair-end sequencing (Read1, 37 bp; Read2, 38 bp).

### CUT&Tag data analysis

Approximately 20 million paired-end reads per sample were sequenced, which is sufficient for robust peak calls with CUT&Tag. Reads were processed for peak calling according to ENCODE ChIP-seq guidelines (https://github.com/ENCODE-DCC/chip-seq-pipeline2)^75^. Briefly, the trimmed, paired-end raw reads were aligned to human reference genome (hg38) using Bowtie 2 (version 2.2.6)^76^, unmapped and multi-mapped reads were filtered using SAMtools (version 1.7)^48^, the ENCODE hg38 blacklist regions (http://mitra.stanford.edu/kundaje/akundaje/release/blacklists/hg38-human/hg38.blacklist.bed.gz) and duplicate reads were filtered using Picard (version 1.126)^77^, high-quality peaks were identified with MACS2 (v.2.2.7.1) with Irreproducible Discovery Rate (IDR) < 0.05. A merged peak list from three biological replicates for each clone was generated and a “concensus peak list”^78^ from two individual clones of the same genotype was produced by filtering out potential random pA-Tn5 enzyme insertion sites^27^. All heatmaps were plotted using deepTools (version 3.5.1)^79^. Differential binding sites (DBS) were detected using the R/Bioconductor package Linnorm (2.20.0)^41^ for CUT&Tag data, the DBS tracks were visualized in Integrative Genomics Viewer (IGV). All experiments were conducted in three biological replicates from each individual clone, and 2 individual clones for each genotype.

### Single-cell RNA sequencing

We generated a high viability (>90%) single cell suspension from three organoids per sample dissociated using papain digestion (Worthington, LK003160). Briefly, organoids were incubated for 30 min at 37 °C in Earle’s Balanced Salt Solution (EBSS)/Albumin-ovomucoid inhibitor/papain with gentle shaking and mechanical dissociation. The pellet was resuspended and incubated with (EBSS)/Albumin-ovomucoid inhibitor/DNAse for 2 min followed by incubation with ovomucoid for 2 min. The cells were washed and resuspended with DMEM/F12. After determination of high cell viability (>90%), the single cell suspension was sequenced on the 10x Genomics platform at the Columbia Genome Center to recover ∼5000 sequenced cells per sample with more than 50,000 reads per cell. The Cell Ranger (5.0.1) pipeline (10x Genomics) was used to align and aggregate all reads from scRNA-seq to the GRCh38 human reference genome with default settings and produce a balanced combined “filtered feature bc matrix” for all samples. The data was imported into the SCANPY software (Version 1.7.1), where genotype information was added. Quality control was conducted to ensure all samples analyzed contained highly consistent quality. Cells expressing a minimum of 200 genes were kept, and counts were normalized for each cell by the total expression, multiplied by 10^6^ and log-transformed. LIGER (Version 2.0.1) software^80^ was used to integrate the different samples and variation in the cells’ transcriptional profile was visualized by uniform manifold approximation and projection for dimension reduction (UMAP) function in SCANPY. To annotate the cell types in brain organoids, we downloaded the gene-cell count matrix and cell type annotation files of the scRNA-seq analysis of human embryonic prefrontal cortex tissue from the cell browser(https://cells.ucsc.edu/). We then utilized this dataset to train the SingleR software (Version 1.0.6 to infer the cell types in our dataset. Differentially expressed genes (DEGs) were determined using model-based analysis of single-cell transcriptomics (MAST, Version 1.12.0) algorithm implemented in Seurat (Version 4.2).

To assess developmental trajectories, we employed two approaches. First, we used Monocle3^81^ to assess pseudotime trajectories of the lineages in our scRNA-seq dataset. Ridgeline plots of cell distribution along pseudotime and Kolmogorav-Smirnov (K-S) test were performed to assess if the distributions were significantly different between mutant and WT organoids. Second, we used the scVelo tool^82^ to analyze RNA velocity of individual cells in WT and mutant organoid datasets. Further analyses of absorption probability and lineage “driver” genes were conducted using the CellRank tool^43^.

### Interval-based enrichment analysis of SETD1A peaks

SETD1A peaks in total and in different genomic regions were subjected to a permutation test implemented in RegioneR^83^ package in R to determine if there is any significant enrichment of these elements within genomic signals identified in previous GWAS studies of SCZ.

### Gene-based enrichment analysis of SETD1A target genes

For the gene-based enrichment analysis, SETD1A peaks were first annotated to known gene targets in the human genome using HOMER^84^, with intergenic SETD1A peaks annotated to their target genes using GREAT^85^. Enrichment among the human SETD1A target gene set of genes implicated in psychiatric disorders (SCZ, BPD, MDD) and neurodegenerative disorders (Parkinson’s disease and Alzheimer’s disease) was determined using DNENRICH^86^, a statistical software package for calculating permutation-based significance of gene set enrichment among genes hit by *de novo* mutations accounting for potential confounding factors such as gene sizes and local trinucleotide contexts. DNENRICH analyses were performed with 100,000 permutations using the following input files: 1) gene name alias file: the default dataset included in the DNENRICH package, 2) gene size matrix file: the default dataset included in the DNENRICH package (for RefSeq genes), 3) gene set file: different sets of high-confidence SETD1A target genes (HGNC symbols available); all genes were equally weighted, 4) mutation list files: lists of genes of interest implicated in psychiatric and neurodegenerative disorders.

### GO term enrichment and motif analysis

Analysis of gene ontology (GO) terms enriched among SETD1A target genes was performed using ToppGene^87^ with default parameters for three categories (Molecular Function, Biological Process and Cellular Component of GO terms). *De novo* motif discovery analysis was performed with HOMER v4.11.1 using automatically generated background (sequences randomly selected from the genome, matched for GC% content).

### WGCNA analysis

For WGCNA analysis of WT organoids at DIV70, we converted the scRNA-seq dataset into Seurat format, subset cells in the EN lineage (including “RG”, “vRG”, “IPC-div”, “IPC-nEN”, “nEN”, “EN” cell types) were analyzed with the hdWGCNA pipeline^88^ with default settings and top 2000 highly variable genes were used for the analysis. After the WGCNA modules were identified in WT, we derived harmonized module eigengenes (hMEs) for both mutant and WT and compared distribution of hMEs of each module in each cell type in the EN lineage using both student t-test and wilcox test implemented in R. Finally, we annotated the potential function of each module using ToppGene (https://toppgene.cchmc.org/).

### Transcriptional component identification

We obtained microarray expression data from the Allen Human Brain Atlas (AHBA) (https://human.brain-map.org/), which provides genome-wide expression data from six adult donors (ages 24–57). In conjunction with HCP-MMP1.0 parcellation images aligned to the native MRI space of each donor’s brain, we used Abagen^89^, and followed the parameter settings established by Dear *et al.*^51^ with one modification: rather than retaining genes based on 50% differential stability^90^, we intersected the 15,946 genes retained after intensity filtering and probe selection with our four gene-of-interest lists—comprising all SETD1A target genes, promoter target genes, intron target genes, and intergenic target genes. To ensure adequate cross-donor representation, we included only regions available in at least three donors, yielding a final dataset of 137 cortical regions. Each gene-of-interest list was intersected with the (137 × 15,946) processed AHBA data matrix, yielding a 137 × N expression matrix (137 cortical regions by N genes, where N corresponds to the number of overlapping genes in each list). These four resultant expression matrices served as the input to the following data analysis. We applied diffusion map embedding (DME) with the same parameters used by Dear *et al*^51^ to each gene list’s expression matrix. For each gene list, we extracted the first five diffusion map components (DC1, DC2, and DC3), which were represented as 137-length vectors, assigning a score to each cortical region. Then we conducted a triplet analysis following the approach outlined by Dear *et al*^51^ for each of the four gene lists. For each gene list, we extracted the first five diffusion map components (DC1-5), which were represented as 137-length vectors, assigning a score to each cortical region. For each component, we aggregated correlation values across the 10 triplet pairs and computed the median absolute correlation as the generalizability score (g), Three components with g>0.5 were selected as generalizable components for downstream analysis. Gene weights were calculated by taking the Pearson correlation between each gene’s expression profile over 137 regions and the component’s regional scores, where a positive and negative correlation produces a positive and negative weight, respectively.

### Correlations of transcriptional components and neuroimaging and macroscale Maps

We followed the correlation framework by Dear *et al*^51^ to examine the relationship between the transcriptional components and nine neuroimaging and macroscale cortical maps with a focus on absolute correlation values instead of hierarchical clustering, allowing us to directly compare the strength of associations between transcriptional components and neuroimaging-derived cortical features across different gene sets.

### Developmental trajectory analysis using BrainSpan

The developmental trajectories of each decile of DC1–DC3 were computed using the procedure described in Dear *et al*^51^. Briefly, the expression of each gene along the time (post-conception days on the log scale) was fitted with a generalized additive model by the statsmodels Python package with alpha = 1 and 12 3rd-degree basis splines as a smoothing function (df = 12, degree = 3 in the BSplines function). Sex and brain region were included as covariates. The fitted data was then averaged by decile of gene weight for each of DC1-DC3 and all deciles were plotted with x axis as log (post-conception days) and y axis as fitted expression level.

### Spatial associations of disease cortical volumes

Following the methodology of Dear *et al*^51^, we analyzed disorder-related cortical shrinkage maps derived from the BrainChart lifespan neuroimaging dataset, which aggregates over 125,000 MRI scans from healthy individuals and patients across development. A key technical consideration was the difference in atlas parcellation: BrainChart-derived shrinkage maps were provided in the Desikan–Killiany (DK) atlas (68 cortical regions per hemisphere), while transcriptional components were computed in the HCP-MMP atlas (180 cortical regions per hemisphere). To align these frameworks, we projected AHBA-derived components from HCP-MMP onto FreeSurfer’s 41k fsaverage surface and re-averaged them to match DK regions, following Dear *et al*^51^. This enabled direct comparisons between the transcriptional components and disorder-related cortical shrinkage values.

For each gene list, we computed Pearson correlations between the three transcriptional components (DC1, DC2, DC3) and cortical shrinkage patterns in ASD, MDD, and SCZ across DK regions. Statistical significance was assessed using spin permutation tests (5,000 permutations) following the Cornblath method^91^, with false discovery rate (FDR) correction applied for multiple comparisons. By conducting this analysis separately for each gene list, we evaluated whether specific subsets of genes drive the observed spatial associations between transcriptional gradients and disorder-related cortical atrophy.

## Data and Code availability

We have uploaded the RNA-seq, CUT&Tag and scRNA-seq data generated in this study in NCBI’s Gene Expression Omnibus under accession code GSE237165, GSE237297 and GSE239864, respectively. There were no unique code or algorithms in this study. All methods used for the analyses are described in the main text and Methods.

## Extended Data Figures

**Extended Data Fig 1.**
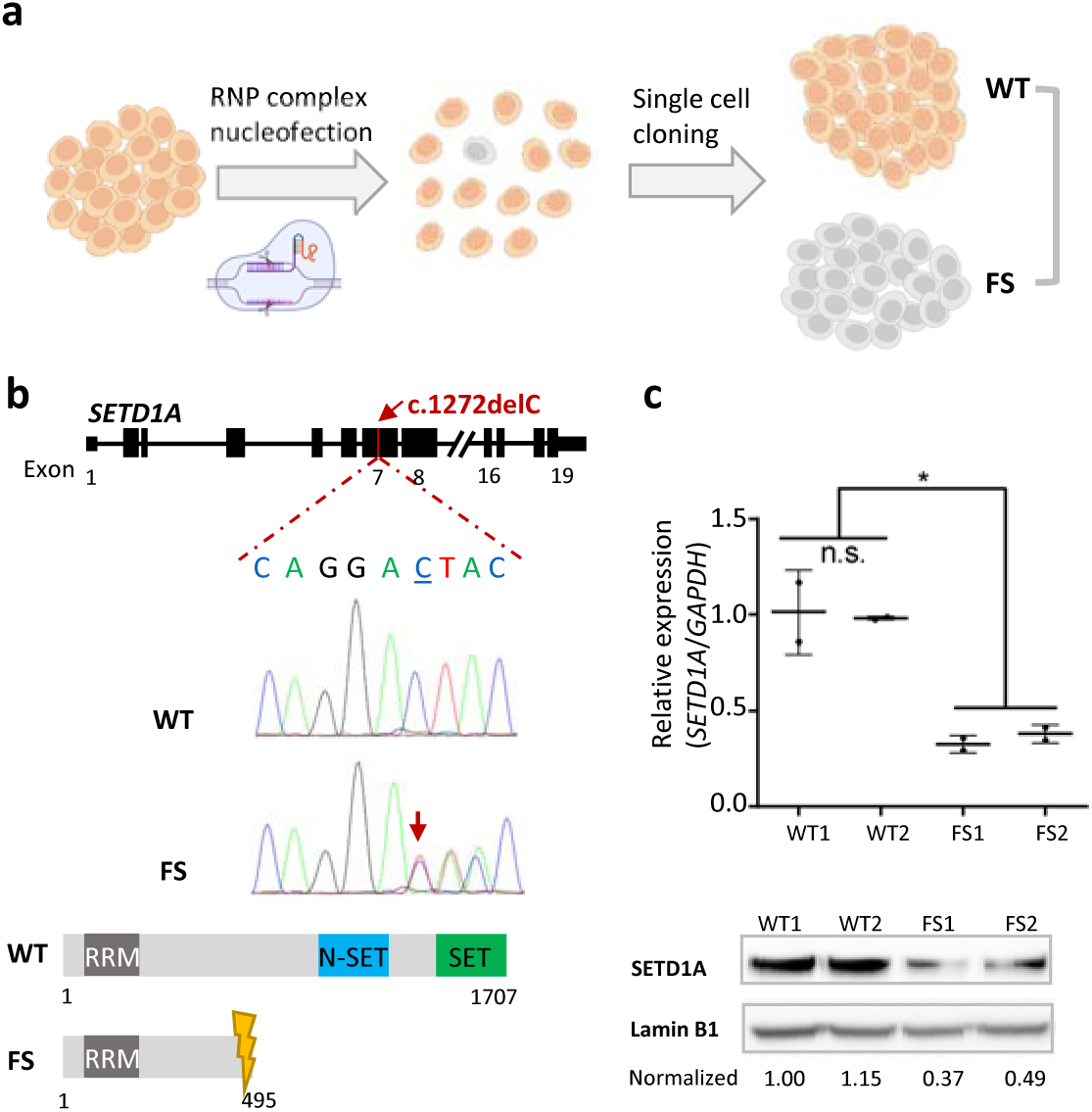
Generation of hiPSC lines harboring a *SETD1A* LoF mutation. **a**. Schematic illustrating the CRISPR/Cas9-mediated generation of the hiPSC lines employed in this study. WT denotes wild-type, FS denotes a frame-shift mutation (c.1272delC) introducing a premature stop codon. RNP: Ribonucleoprotein. **b,** Representative chromatograms of the SETD1A alleles in WT and FS hiPSC lines. Depicted are the genomic structure of the SETD1A gene as well as the full length (1707 aa) WT and predicted truncated (495 aa) mutant SETD1A protein, and key functional domains (RRM: RNA recognition motif, SET: Suppressor of variegation 3–9, enhancer of zeste and trithorax, N-SET: N-terminal of SET). **c,** Results of quantitative RT-qPCR (top) and western blot analysis (bottom) of SETD1A expression in the four WT and mutant hiPSC clones employed in this study revealing reduction in SETD1A mRNA and protein levels in mutant clones. TaqMan primers spanning exon 3-4 of SETD1A mRNA were used in RT-qPCR assay. SETD1A protein levels were probed with mouse monoclonal antibody sc-515590 (Santa Cruz Biotechnology), quantified and normalized to a housekeeping gene encoding Lamin B1, and fold changes were shown as a relative value to WT1. Representative data of 2 independent experiments were shown.

**Extended Data Fig 2.**
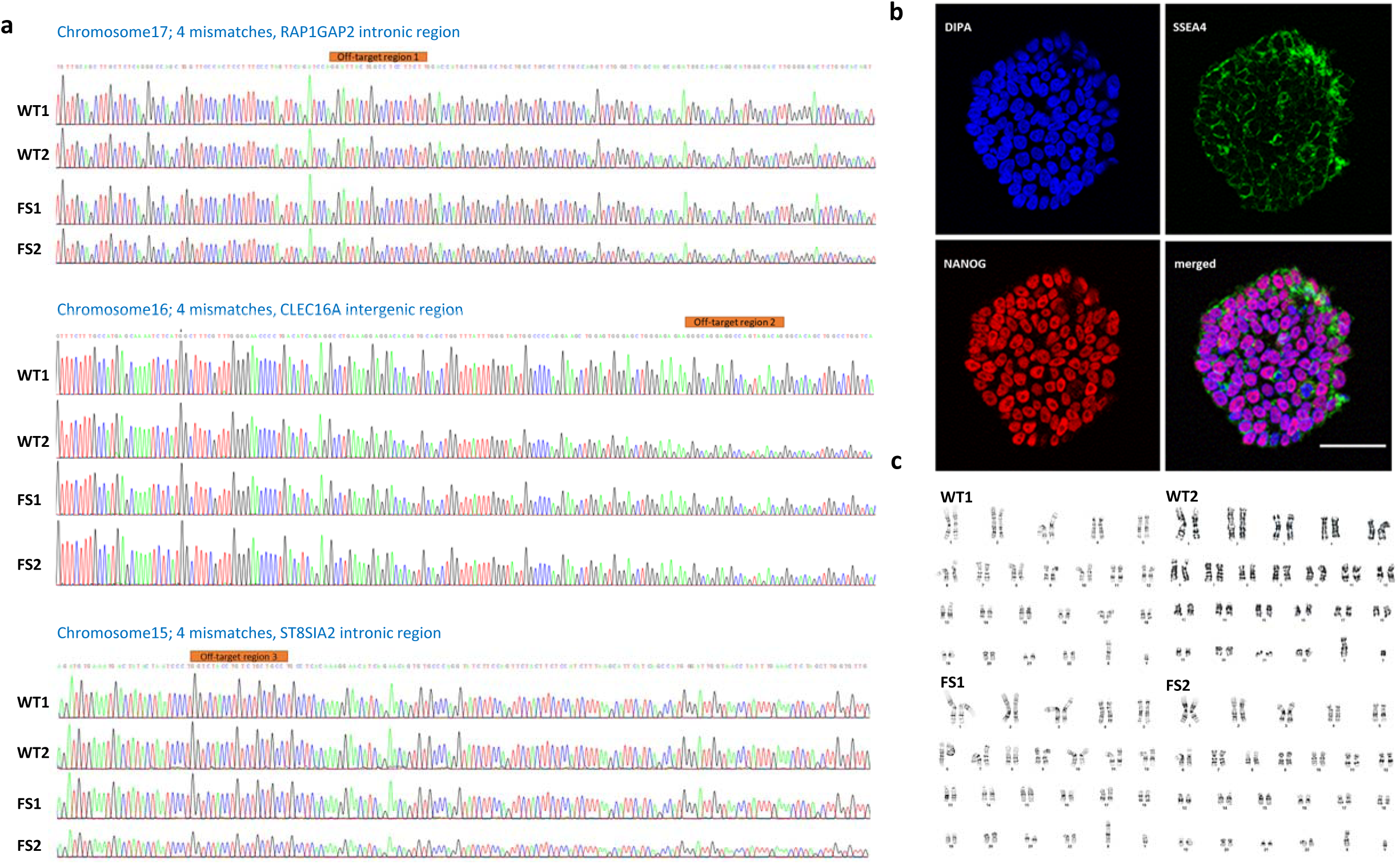
Characterization of hiPSC lines. **a**, Analysis of off-target CRISPR/Cas9 activity in WT and mutant hiPSC lines. The top three predicted off-target regions were PCR amplified and subjected to Sanger sequencing. Traces for all clones show no indels or mismatches in a 500-700 bp region around the predicted off-target regions. **b,** Representative image of immunofluorescence staining of hiPSCs with embryonic stem cell markers SSEA4 and NANOG as well as DAPI in hiPSCs. Scale bar, 50µm. **c,** Karyotyping of WT and mutant hiPSC lines demonstrates normal karyotype distribution in all clones.

**Extended Data Fig 3.**
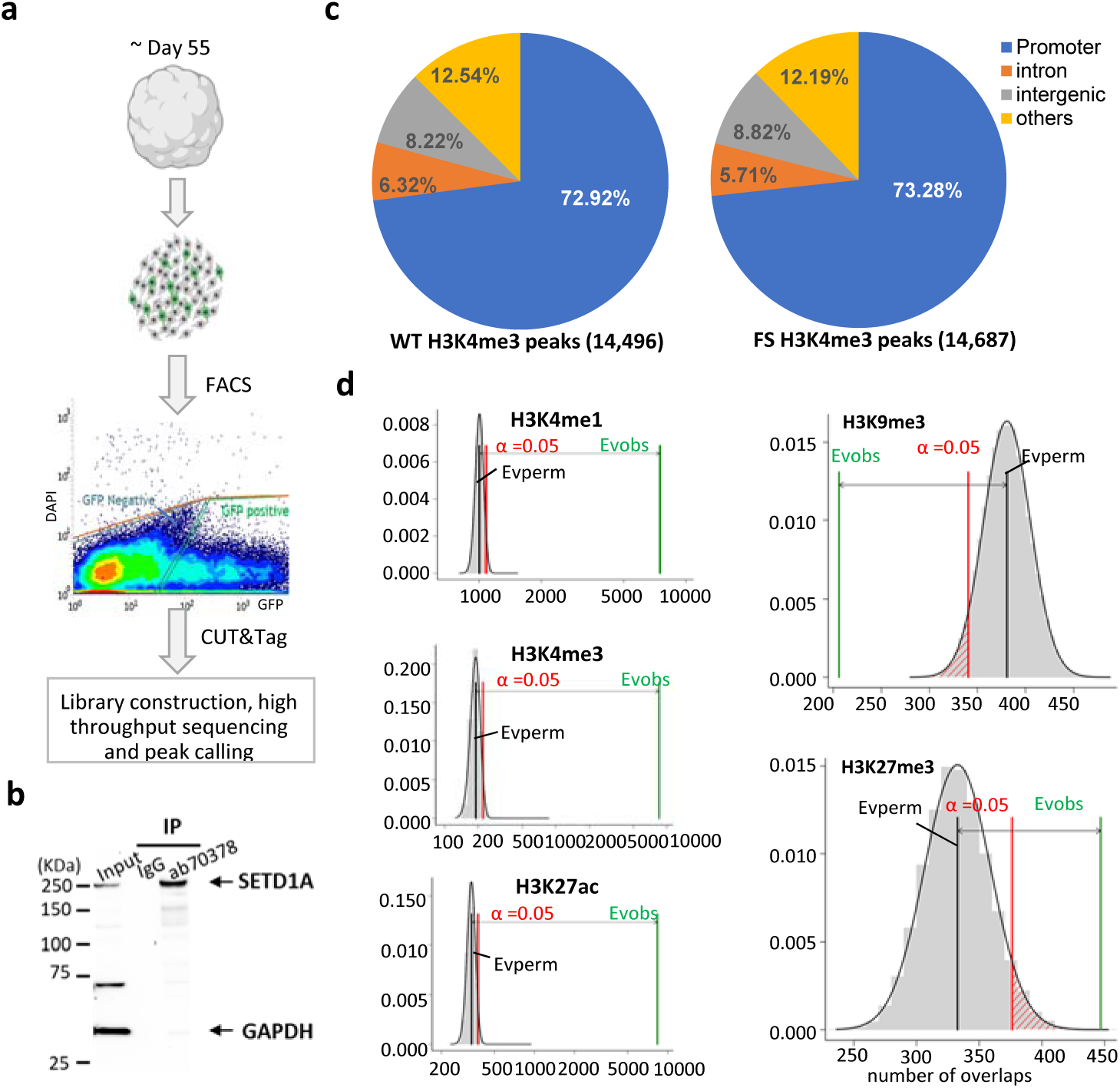
Identification of SETD1A genomic targets and H3K4me3 binding on cortical neurons from WT and mutant organoids. **a**, Schematic outline of the identification of SETD1A genomic targets using CUT&Tag on cortical neurons derived from WT and mutant DIV70 organoids transduced with AAV1-hSYN-eGFP virus via FACS. The output sequence data were analyzed using the well-established ENCODE chip-seq pipeline2 for read alignment, quality control and peak calling. **b,** Validation of the specificity of SETD1A antibody employed in the current study by Immunoprecipitation (IP) in HEK293T cells. **c,** Genomic distribution of H3K4me3 peaks in WT (n = 14,496, top) and FS (n = 14,687, bottom) neurons. **d,** Permutation tests (n = 100,000, using circular Randomize Regions in RegioneR) indicate a higher and lower than expected by chance overlap between histone marks and SETD1A peaks and (p = 1×10-5, Z = 140.0 for H3K4m1; p = 1×10-5, Z = 436.2 for H3K4m3; p = 1×10-5, Z = 332.7 for H3K27ac; p = 1×10-5, Z = −7.2 for H3K9me3; p = 1×10-5, Z = 4.3 for H3K27me3). x axis depicts the number of overlaps and y axis depicts the density of expected number of overlaps as determined by permutation. Evperm denotes the expected average number of overlaps; Evobs denotes the observed number of overlaps. Permutation P value threshold is set at 0.05.

**Extended Data Fig 4.**
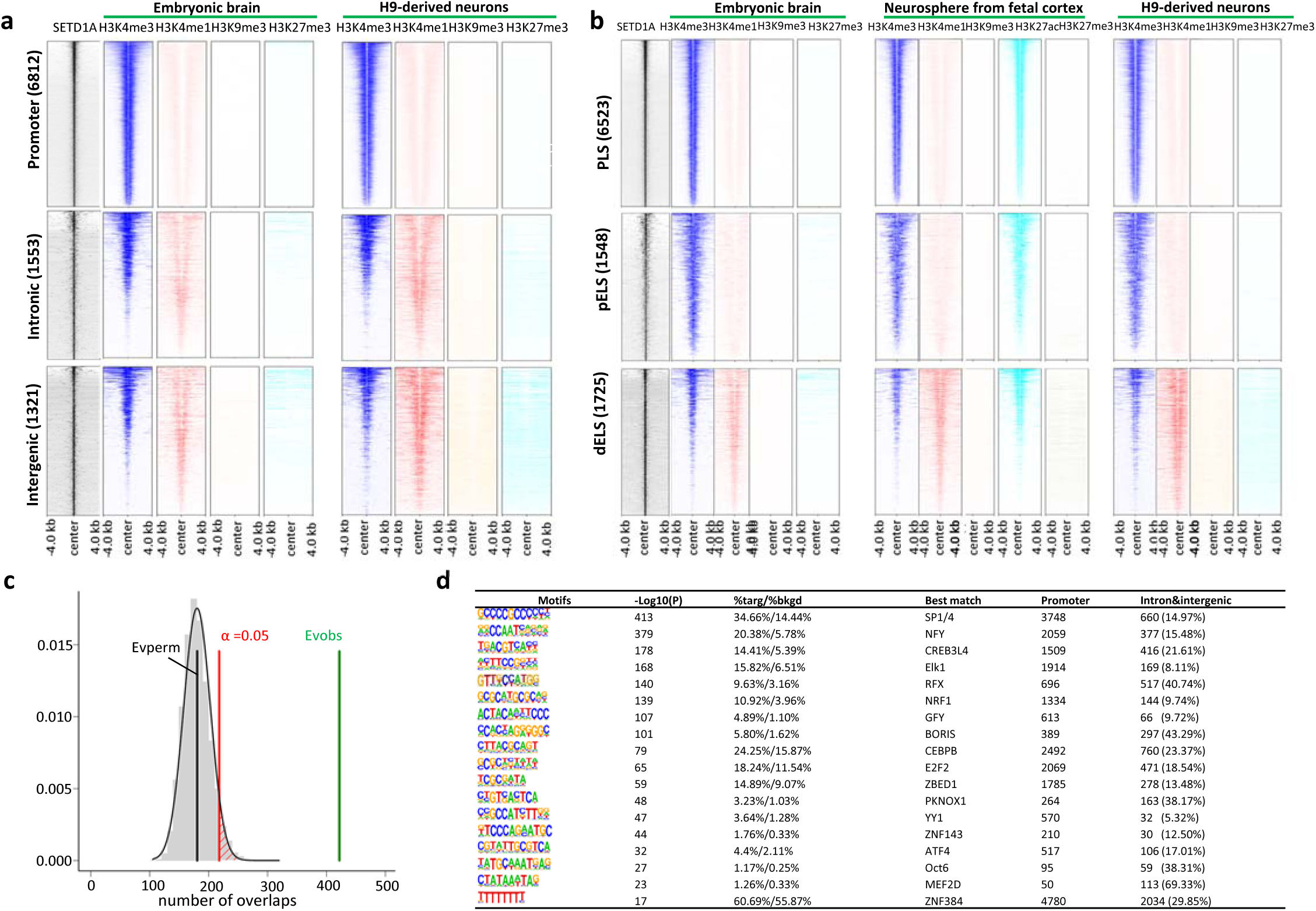
Histone marks, higher-order chromatin contacts and TF motifs of SETD1A binding sites. **a,** Heatmap of histone marks centered on SETD1A peaks (n = 10218) across a ±4-kb window. Histone modification profiles were extracted from ENCODE datasets from the human embryonic brain and H9-derived neurons and compared to the SETD1A genomic occupancy map. SETD1A peaks were clustered in proximal promoters (<1kb) (top, 6812 peaks), intronic (middle, 1553 peaks) and intergenic regions (bottom, 1321 peaks). **b,** Heatmap of histone marks centered on SETD1A peaks (n = 10218) across a ±4 kb window. PLS: promoter-like sites, pELS: proximal enhancer-like sites, dELS: distal enhancer-like sites. **c,** Permutation tests (n = 100,000, using circular Randomize Regions in RegioneR) between HiC loci identified in the cortical and subcortical plate of human fetal cerebral cortex and SETD1A peaks. x axis depicts the number of overlaps and y axis depicts the density of expected number of overlaps determined by permutation. **d,** *De novo* TF motifs enriched in SETD1A peaks (n = 10218).

**Extended Data Fig 5.**
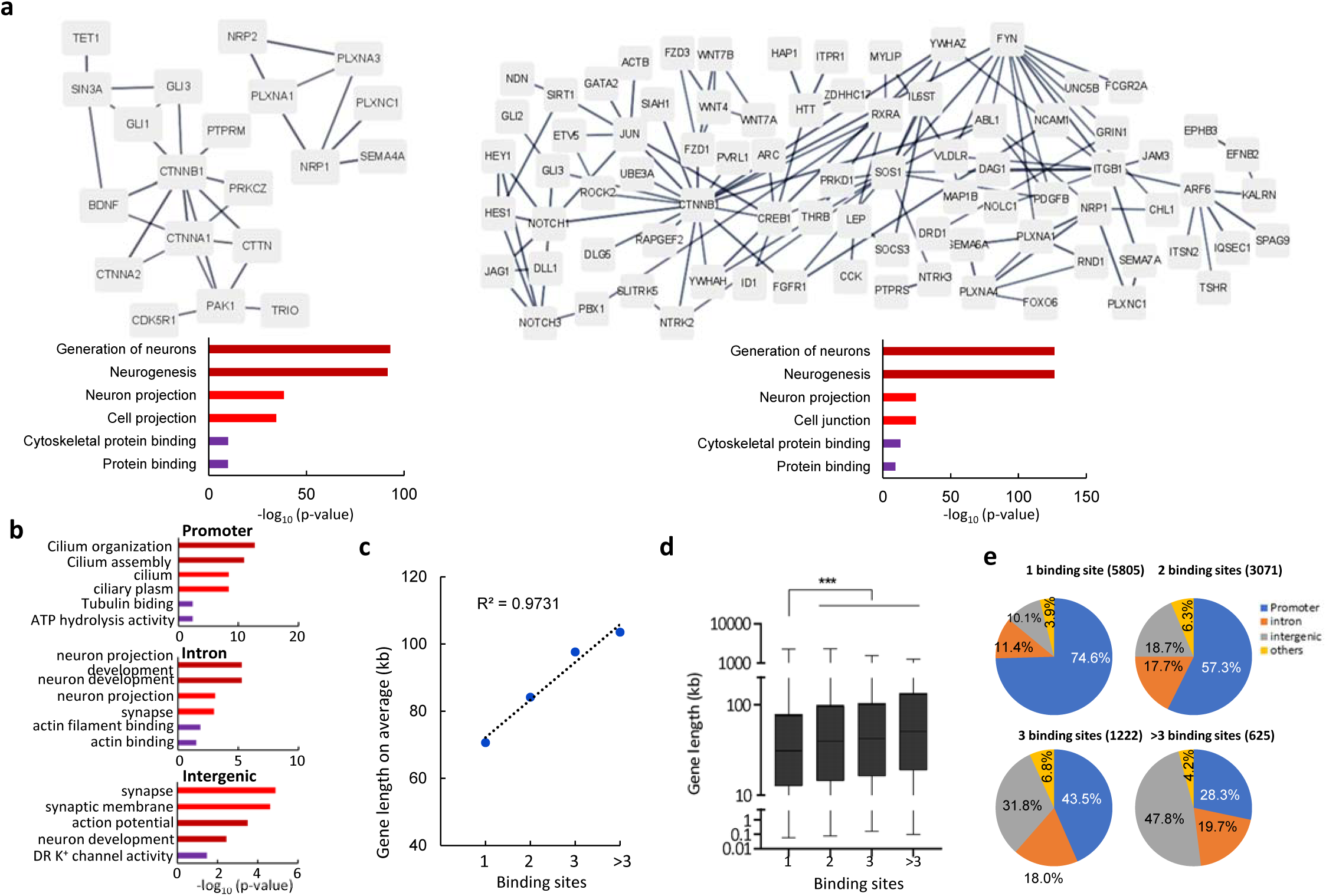
Functional enrichment and features of SETD1A target genes. **a,** PPI networks analysis for intronic-(upper left) and intergenic-(upper right) bound SETD1A target genes associated with the “neuron development” GO term. The interaction score is 0.9. GO term enrichment analysis for target genes with SETD1A bound intronic (bottom left) or intergenic regions (bottom right). **b,** GO term enrichment analysis for SETD1A binding sites with an RFX3 motif in promoter (top), intronic (middle) and intergenic regions (bottom). **c,** Correlation between the average target gene length (RefSeq transcription start to termination site) and the number of SETD1A binding sites. **d,** Boxplot of gene lengths for target genes with different numbers of SETD1A binding sites. * p < 0.05, ** p < 0.01, *** p < 0.001, unpaired Student’s t test. **e,** Genomic distribution of SETD1A peaks among target genes classified by the number of SETD1A binding sites in their regulatory regions.

**Extended Data Fig 6.**
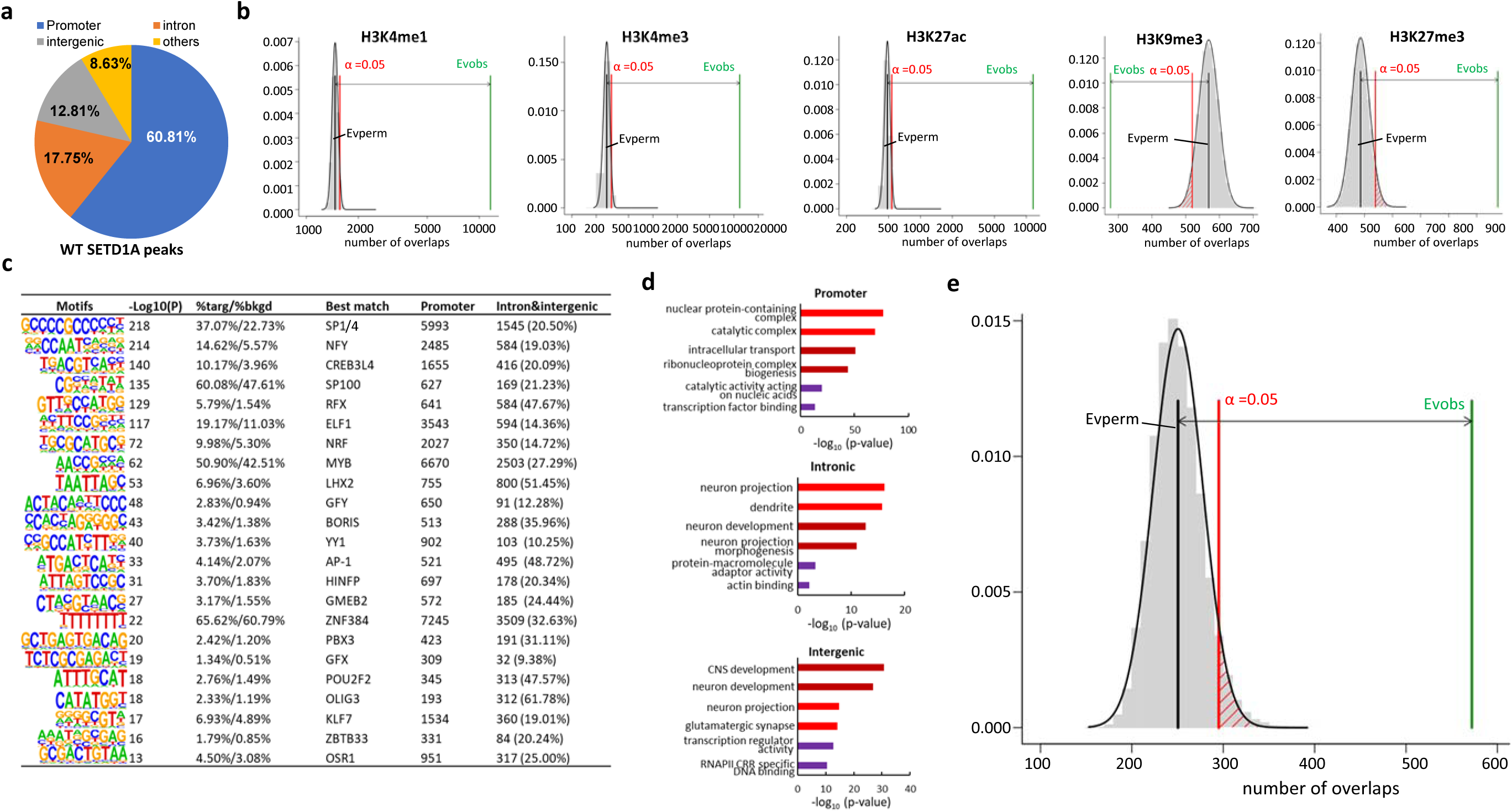
Genome-wide characterization of SETD1A binding sites in an independent batch of organoids. **a**, Genomic distribution of SETD1A peaks in WT neurons. **b**, Permutation tests (n = 100,000, using circular Randomize Regions in RegioneR) indicate a higher than expected by chance overlap between histone marks and SETD1A peaks. x axis depicts the number of overlaps and y axis depicts the density of expected number of overlaps as determined by permutation. EVperm denotes the expected average number of overlaps; EVobs denotes the observed number of overlaps. Permutation *P* value threshold is set at 0.05. **c**, *De novo* TF motifs enriched in SETD1A peaks. **d**, GO analysis of genes with SETD1A binding sites in promoter, intronic and intergenic regions indicates that genes targeted by SETD1A at intronic and intergenic enhancer sites are more likely involved in neuronal-specific processes. **e**, Permutation tests (n = 100,000, using circular Randomize Regions in RegioneR) between HiC loci identified in the cortical and subcortical plate of human fetal cerebral cortex and SETD1A peaks. x axis depicts the number of overlaps and y axis depicts the density of expected number of overlaps determined by permutation.

**Extended Data Fig 7.**
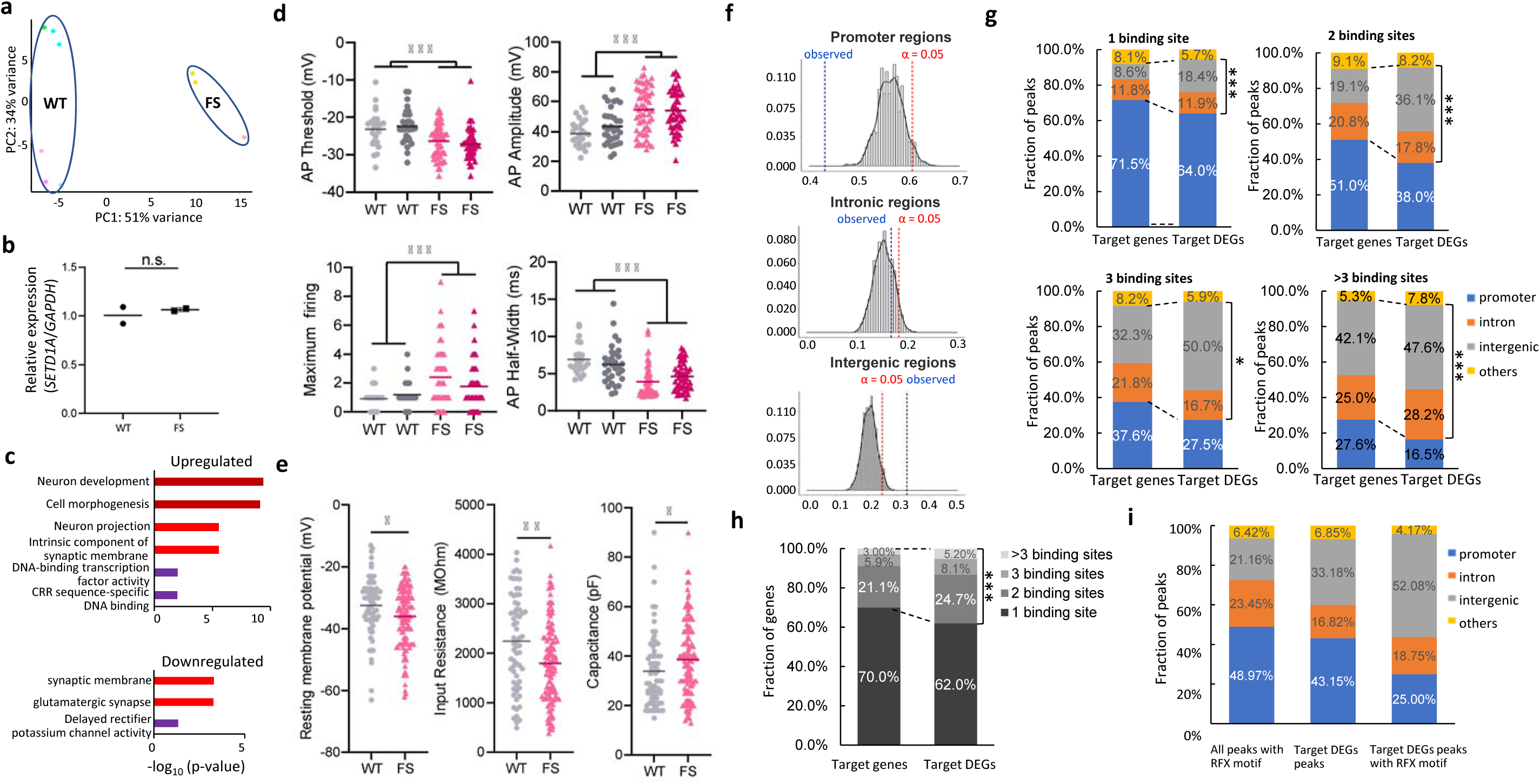
Transcriptional alterations induced by *SETD1A* LoF mutation in relation to SETD1A genomic binding sites and impact on electrophysiological properties. **a**, PCA analysis of bulk RNA-seq data from neurons sorted from WT and mutant organoids. **b**, Results of quantitative RT-qPCR in neurons sorted from WT and mutant organoids. TaqMan primers spanning exon 3-4 of SETD1A mRNA are used in RT-qPCR. **c**, GO term enrichment analysis of upregulated (upper) and downregulated (bottom) SETD1A target DEGs. **d**, Quantification of action potential (AP) threshold (top left), AP amplitude (top right), Maximum firing (bottom left) and AP half width (bottom right) in neurons derived from two independent WT and mutant iPSC lines **e**, Quantification of resting membrane potential (left), input resistance (middle) and capacitance (right). **f**, Permutation plots of SETD1A peaks overlapping with DEGs located in promoter (top), intronic regions (middle) and intergenic (middle). X axis depicts the percent of SETD1A peaks in tested regions, and y axis depicts the density of SETD1A peaks calculated by permutation. “observed” denotes the actual observed number of overlaps. Permutation P value threshold is set at 0.05. **g**, Comparison of the genomic distribution of SETD1A peaks between target DEGs and all SETD1A target genes classified by the number of SETD1A binding sites in their regulatory regions. * p < 0.05, ** p < 0.01, *** p < 0.001, chi square test. **h**, Percent of genes carrying different number of SETD1A binding sites. **i**, Genomic distribution of SETD1A peaks carrying an RFX motif (left), overlapping with DEGs (middle), and overlapping with DEGs and also carrying RFX motif (right), respectively.

**Extended Data Fig 8.**
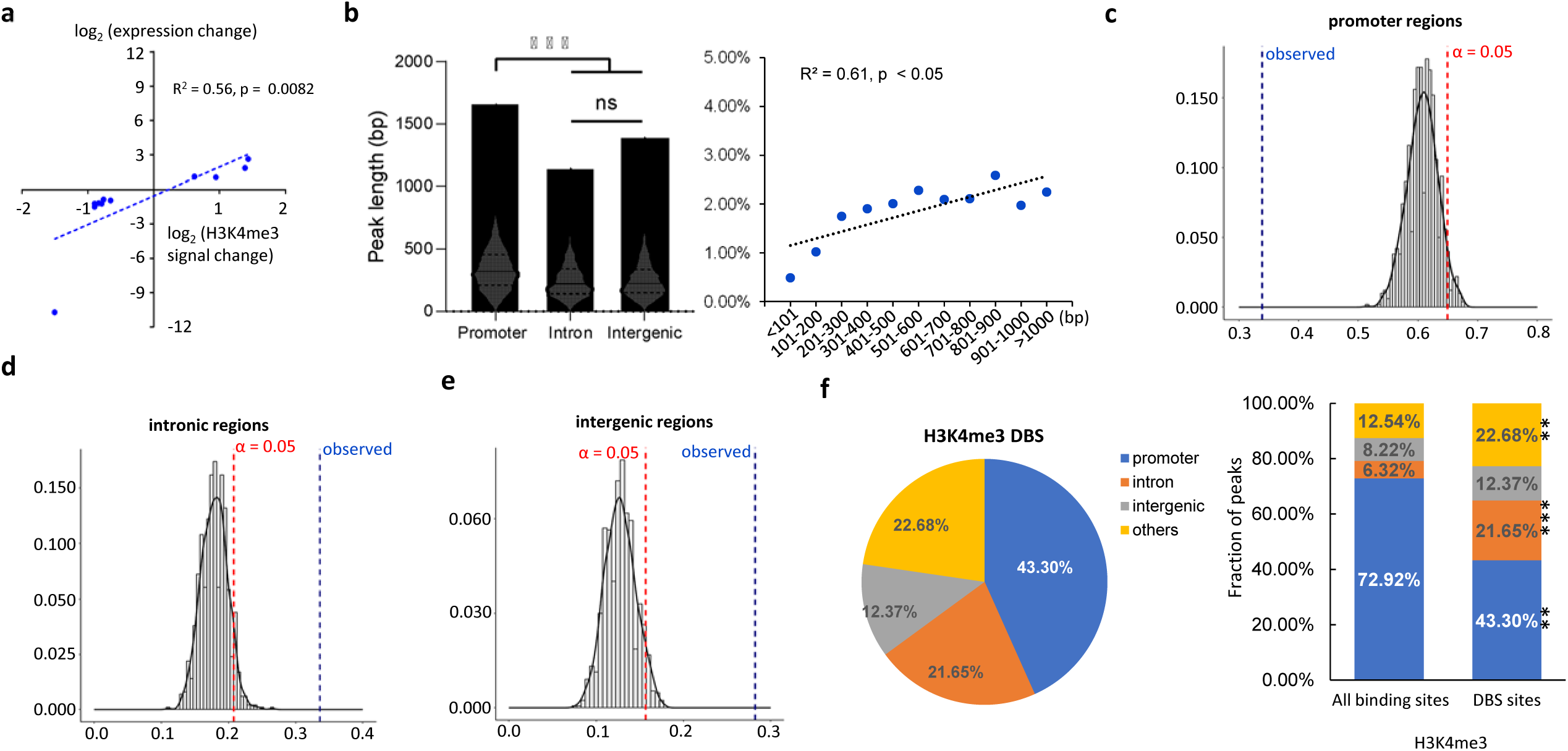
Characterization of SETD1A DBS and differential H3K4me3 signal. **a**, Correlation analysis between SETD1A DBS, differential H3K4me3 signal and DEGs (n = 11) found a positive correlation between the direction of DEG expression and SETD1A/H3K4me3 occupancy changes in associated DBS; x axis depicts log_2_ binding occupancy fold change, y axis depicts log_2_ gene expression fold change. **b**, Analysis depicting the distribution of SETD1A peak lengths across various genomic regions (left) and examination of the correlation between the density of SETD1A DBS and peak lengths (right). **c-e**, Results of permutation tests of SETD1A peaks carrying DBS in promoter, intronic and intergenic regions. x axis depicts the percent of SETD1A peaks carrying DBS in tested regions, and y axis depicts the density of SETD1A peaks calculated by permutation. “Observed” denotes the actual observed number of overlaps. Permutation *P* value threshold is set at 0.05. **f**, Genomic distribution of sites with differential H3K4me3 signal (top) and comparison of the genomic distribution of sites with differential H3K4me3 signal and H3K4me3 binding sites as a whole. * p < 0.05, ** p < 0.01, *** p < 0.001, chi square test.

**Extended Data Fig 9.**
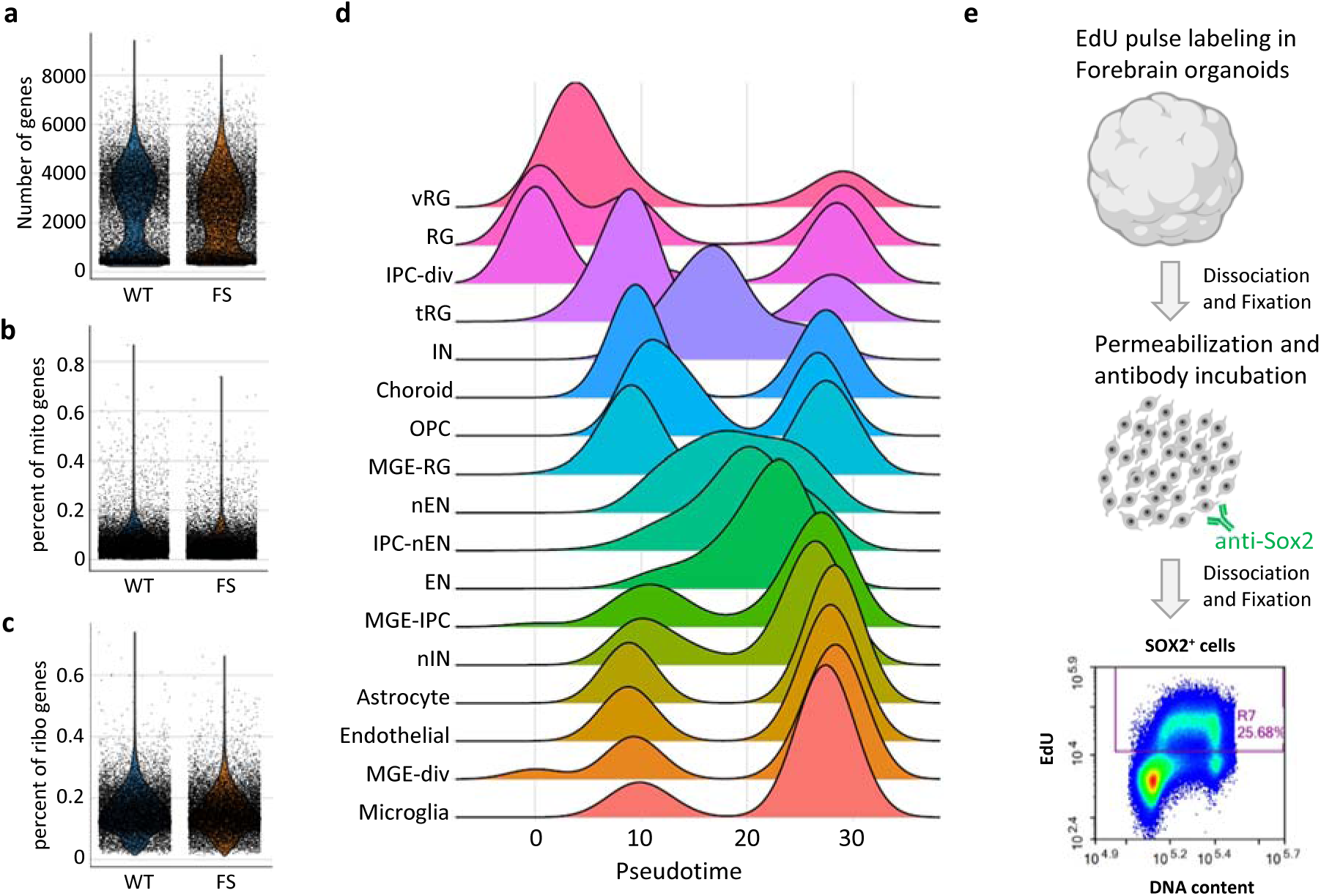
Quality control assessment of scRNA-seq data from DIV70 organoids. **a**, Number of genes identified by scRNA-seq in WT and mutant organoids at DIV70. **b**, Percent of mitochondrial genes identified by scRNA-seq in WT and mutant organoids at DIV70. **c**, Percent of ribosomal genes identified by scRNA-seq in WT and mutant organoids at DIV70. **d**, Plots depicting cellular identities along the pseudotime in forebrain organoids at DIV70. **e**, Schematic of FACS analysis to quantitatively determine the proliferation and neurogenesis rate *via* flow cytometry.

**Extended Data Fig 10.**
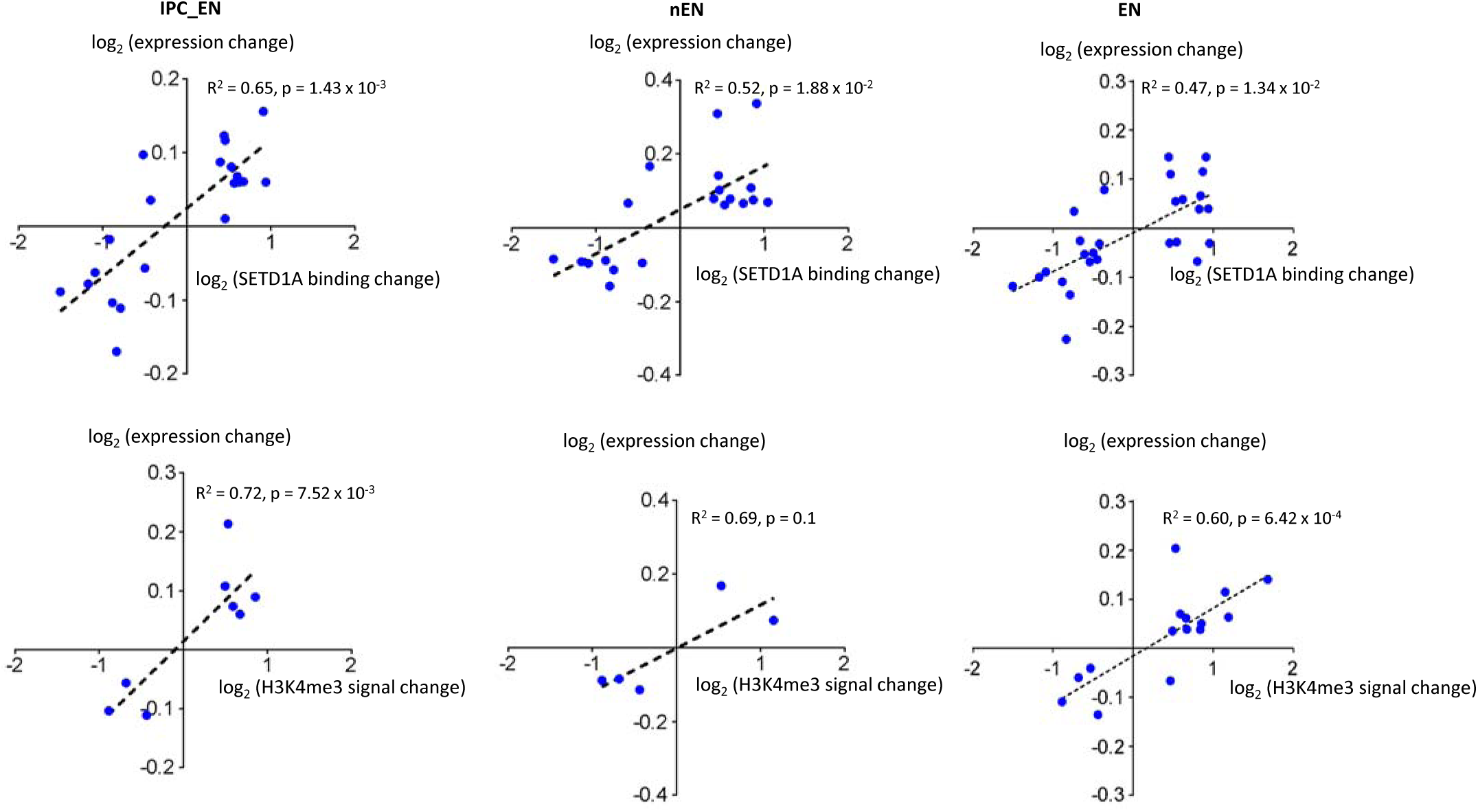
Correlation analysis between SETD1A DBS and EN lineage DEGs. Correlation analysis between SETD1A DBS and DEGs in IPC_EN (upper left), nEN (upper middle), and EN (upper right) revealed a positive association between the direction of differential gene expression and changes in SETD1A occupancy at associated DBS. Similarly, analysis of SETD1A DBS and DEGs in IPC_EN (bottom left), nEN (bottom middle), and EN (bottom right) demonstrated a positive correlation between DEG expression direction and alterations in H3K4me3 levels. In both analyses, the x-axis represents the log₂ fold change in binding occupancy, while the y-axis represents the log₂ fold change in gene expression.

**Extended Data Fig 11.**
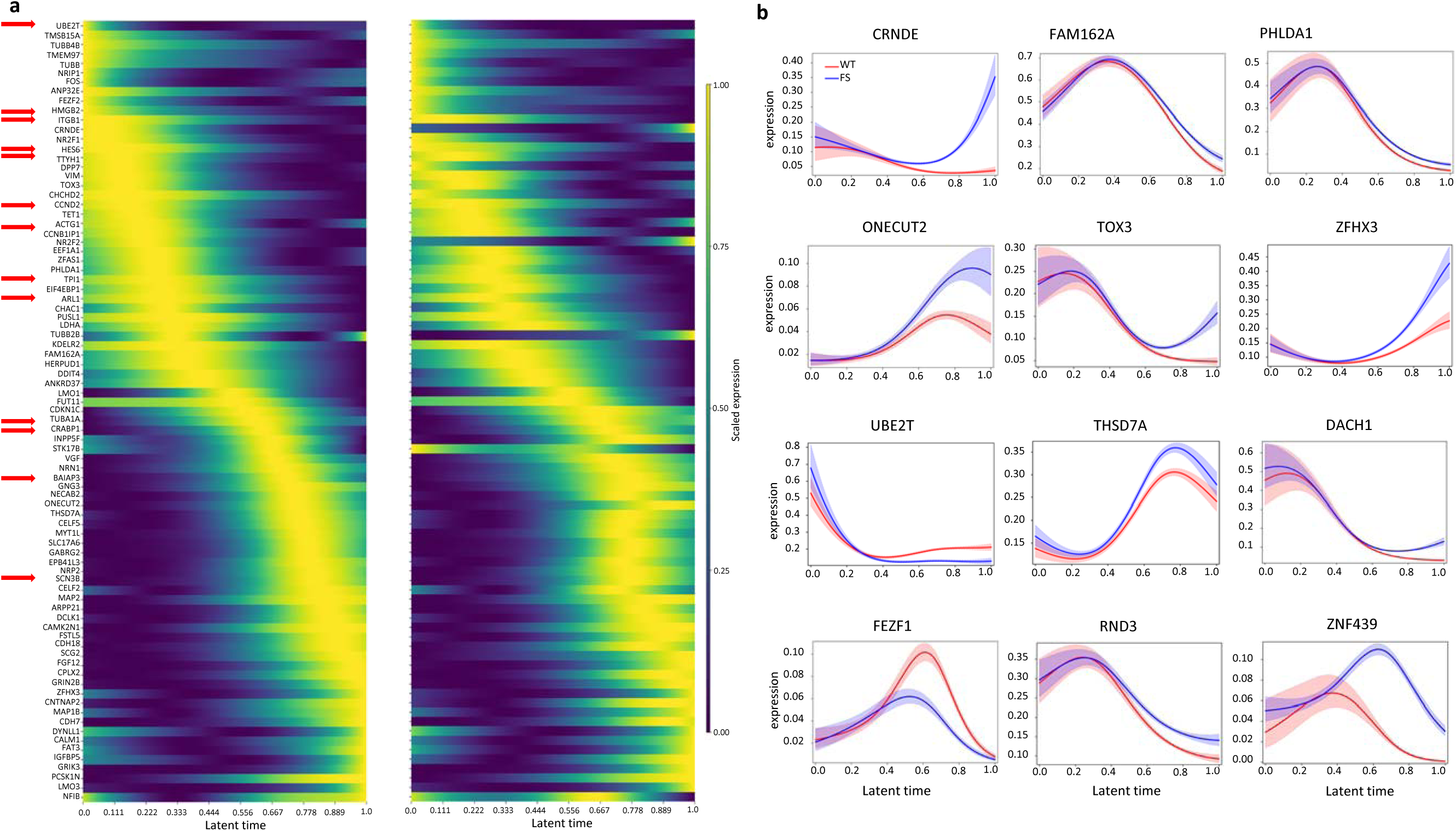
Latent time analysis of dysregulated EN lineage driver genes. **a**, Heatmap of 83 SETD1A target EN lineage driver genes over latent time. b, Latent time plots along velocity latent time of 12 high confidence representative examples of SETD1A target and non-target dysregulated EN lineage driver genes that display consistent changes in both scRNA- and bulk RNA-seq EN datasets. Genes listed on the top two rows, UBE2T and THSD7A are SETD1A target genes.

**Extended Data Fig 12.**
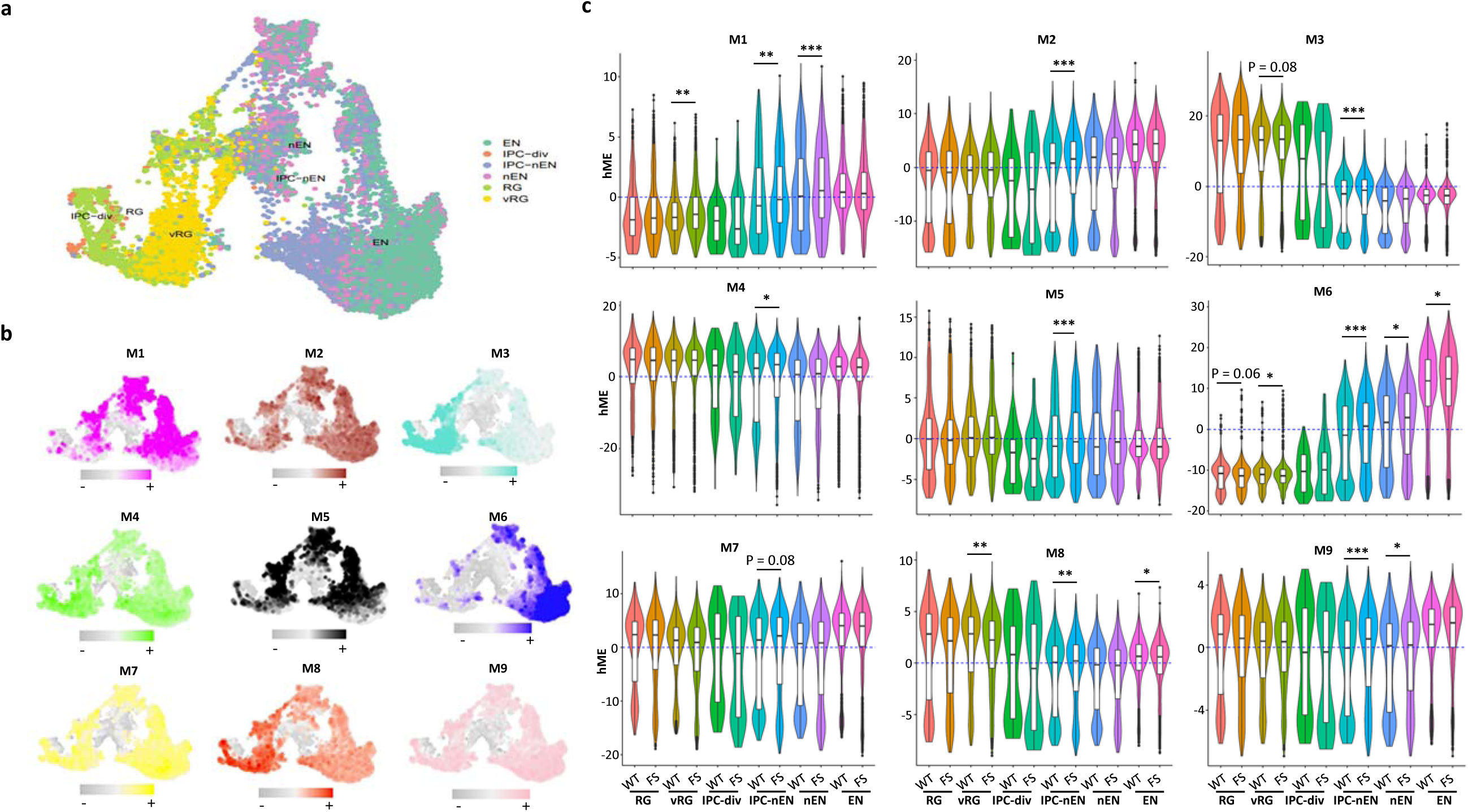
Distribution of WGCNA modules and expression of module genes across different cell types. **a-b**, Distribution of WGCNA modules across different cell types identified in DIV70 organoids. **c**, Expression patterns of module genes across different cell types in WT and mutant organoids. hME: harmonized module eigengenes. * p < 0.05, ** p < 0.01, *** p < 0.001, wilcox test.

**Extended Data Fig 13.**
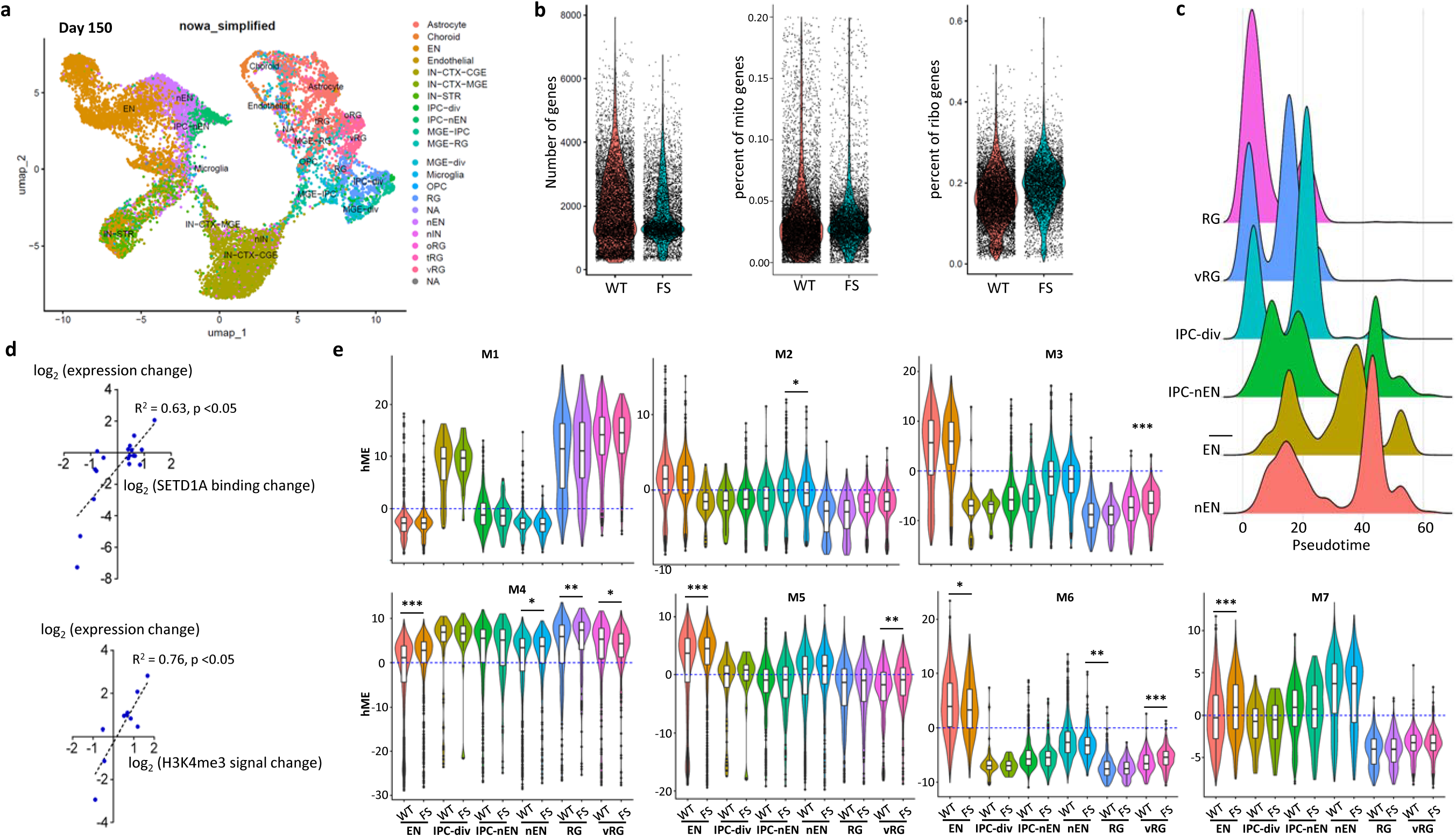
Asynchronous developmental trajectories in DIV150 forebrain organoids. **a,** Cell type annotation, based on human fetal brain references, revealed that most cell types present in the developing fetal cortex are identifiable in DIV150 organoids. The UMAP visualization, color-coded by annotated cell type, underscores the fidelity of these organoids in recapitulating human neurodevelopment. **b,** Number of genes (left), percent of mitochondrial genes (middle) and percent of ribosomal genes (right) identified by scRNA-seq in WT and mutant organoids at DIV150. **c,** Plots depicting cellular identities along the pseudotime in forebrain organoids at DIV150. **d,** Correlation analysis between SETD1A DBS and DEGs identified in EN at DIV150 displays a positive correlation between the direction of DEG expression and SETD1A/H3K4me3 signal changes **e,** Expression patterns of module genes across different cell types in WT and mutant organoids. hME: harmonized module eigengenes. * p < 0.05, ** p < 0.01, *** p < 0.001, wilcox test.

**Extended Data Fig 14.**
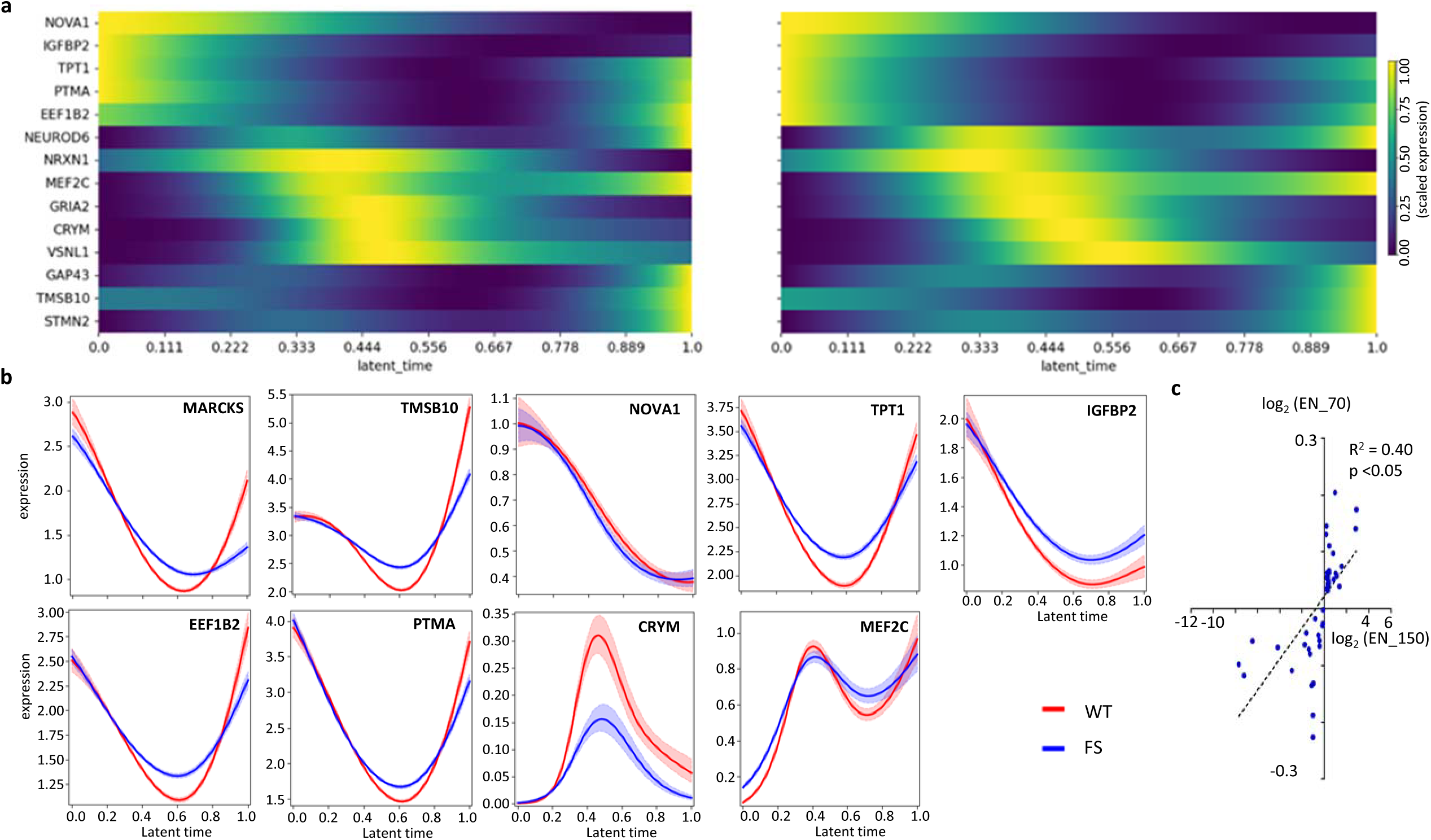
Latent time analysis of dysregulated EN lineage driver genes. **a**, Heatmap of 14 SETD1A target EN lineage driver genes over latent time. **b**, Latent time plots along velocity latent time of 9 representative examples of SETD1A target and non-target dysregulated EN lineage driver genes that exhibit changes in scRNA-seq EN datasets. Genes listed on the top two rows are SETD1A target genes, genes on the last row are non-SETD1A target genes. **c**, Correlation analysis of 49 genes that were consistently dysregulated in both DIV70 and DIV150 in terms of fold-change direction between DIV70 and DIV150 in ENs.

**Extended Data Fig 15.**
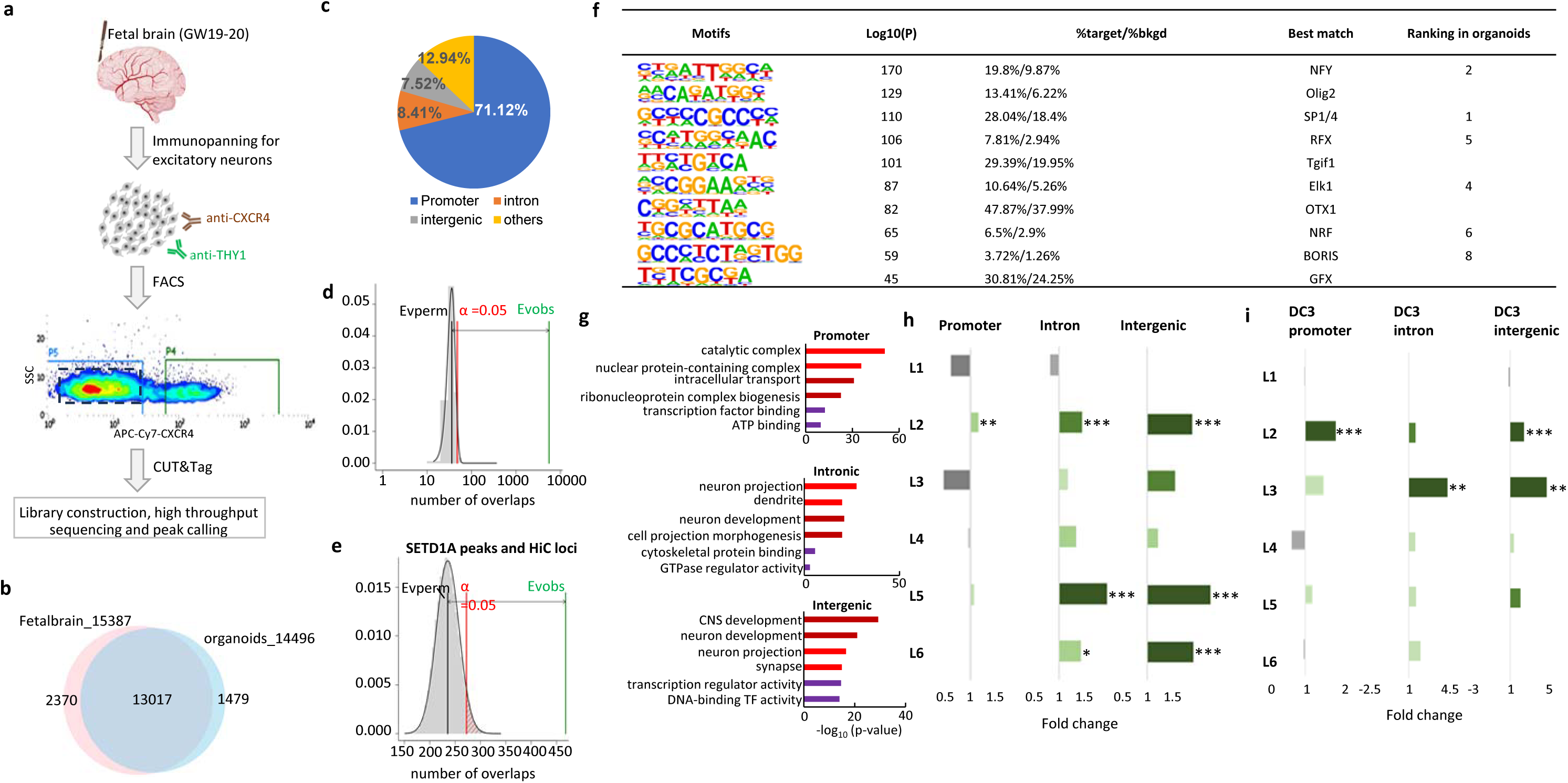
Characterization of SETD1A binding sites and associated genes in EN from human fetal frontal cortices. **a**, Schematic outline of the identification of SETD1A genomic targets using CUT&Tag on EN derived from healthy fetal brain at gestation week 19-20 (GW19-20). **b,** Venn diagram depicting the overlap between SETD1A peaks in EN from fetal cortices (n=15387) and forebrain organoids (n=14496) H3K4me3 peaks. **c,** Genomic distribution of overlapping SETD1A peaks. d, Permutation tests (n = 100,000, using circular Randomize Regions in RegioneR) assessing the overlap (left) between SETD1A peaks in EN from fetal brains (n=13,211) and forebrain organoids (n=10,218). **e,** Permutation tests between HiC loci identified in the cortical and subcortical plate of human fetal cerebral cortex and SETD1A peaks. x axis depicts the number of overlaps and y axis depicts the density of expected number of overlaps determined by permutation. **f,** Top 10 De novo TF motifs enriched in SETD1A peaks (n = 13211). **g,** GO analysis of genes with SETD1A binding sites in promoter, intronic and intergenic regions. **h,** Enrichment of cortical layer-specific marker genes among all SETD1A target genes in promoter, intronic and intergenic regions. **i**, Enrichment of SETD1A target in promoters, intronic regions, and intergenic regions within DC3 among cortical layer-specific marker genes.

**Extended Data Fig 16.**
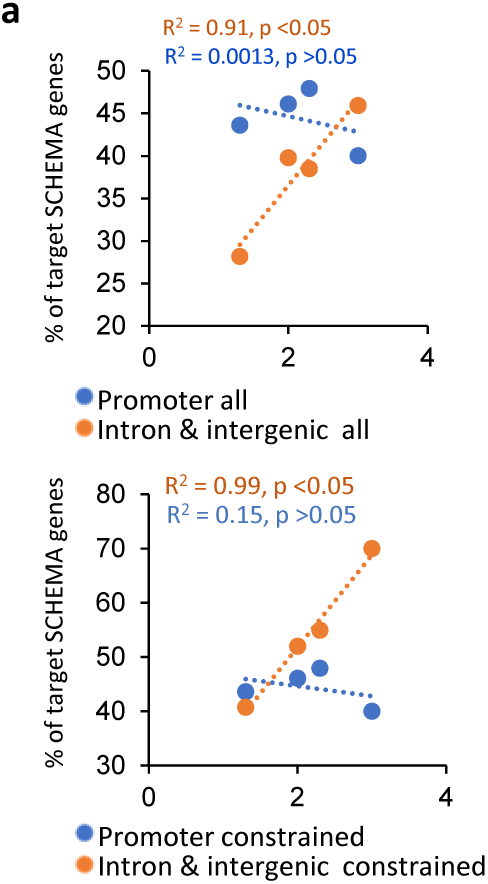
Correlation Between SCHEMA Gene Lists at Varying FDR Thresholds and SETD1A Binding Across Genomic Regions. **a,** Correlation analysis between SCHEMA gene lists at various FDR thresholds (<0.05, <0.01, <0.005, and <0.001) and the percentage of SCHEMA genes—or constrained SCHEMA genes (pLI > 0.9)—harboring SETD1A binding sites across distinct genomic regions.

## Table Legends

**Supplementary Table 1: SETD1A and H3K4me3 Peaks Identified in Sorted Cortical Neurons from WT and Mutant Forebrain Organoids at DIV70**

**Supplementary Table 2: Publicly Available Histone Modification Profiles, GO Analysis and Human-Gained Enhancer Interaction Profiles in Fetal Cerebral Cortices**

**Supplementary Table 3: Motif Analysis of SETD1A Peaks and Functional Categorization of Target Genes**

**Supplementary Table 4: De Novo Motif Analysis and Genomic Annotation of SETD1A Peaks Supplementary Table 5: Differentially Expressed Genes in Mutant Neurons, SETD1A Peak Overlap, and Enriched Motifs**

**Supplementary Table 6: Differential Binding Strength of SETD1A and H3K4me3 Peaks in Sorted Cortical EN and Their Overlap with DEGs**

**Supplementary Table 7: Differential Gene Expression and Driver Gene Identification in Distinct Cell Types of Forebrain Organoids at DIV70**

**Supplementary Table 8: Expression Profiles, GO Analysis, Distribution Comparison of WGCNA Modules in EN Lineage, and Overlap Between DEGs and Module Genes**

**Supplementary Table 9: GO Analysis of DEGs in IPC_nEN, nEN, and ENs at DIV70 and Upregulated Genes Enriched in OXPHOS in IPC_nEN**

**Supplementary Table 10: DEGs and Driver Genes in EN at DIV150 and Overlap Analysis Between DIV150 and DIV70 DEGs**

**Supplementary Table 11: Expression Profiles, GO Analysis, and Overlap of DEGs with WGCNA Modules in DIV150 EN lineage**

**Supplementary Table 12: Analysis of SETD1A and H3K4me3 Peaks in Human Fetal Brain, Genomic Annotations**

**Supplementary Table 13: Gene Lists and GO Enrichment Analysis of SETD1A Target Components (DC1-DC3) and Correlations with Neuroimaging Maps**

**Supplementary Table 14: Cortical Layer Specificity of SETD1A Target Genes within DC1-3 and Overlap with SCZ-Associated Genes**

